# Structural and functional insights into nitrosoglutathione reductase from *Chlamydomonas reinhardtii*

**DOI:** 10.1101/2020.08.31.275875

**Authors:** Andrea Tagliani, Jacopo Rossi, Christophe H. Marchand, Marcello De Mia, Daniele Tedesco, Gurrieri Libero, Maria Meloni, Giuseppe Falini, Paolo Trost, Stéphane D. Lemaire, Simona Fermani, Mirko Zaffagnini

## Abstract

Protein S-nitrosylation plays a fundamental role in cell signaling and nitrosoglutathione (GSNO) is considered as the main nitrosylating signaling molecule. Enzymatic systems controlling GSNO homeostasis are thus crucial to indirectly control the formation of protein S-nitrosothiols. GSNO reductase (GSNOR) is the key enzyme controlling GSNO levels by catalyzing its degradation in the presence of NADH. Here, we found that protein extracts from the microalga *Chlamydomonas reinhardtii* catabolize GSNO *via* two enzymatic systems having specific reliance on NADPH or NADH and different biochemical features. Scoring the Chlamydomonas genome for orthologs of known plant GSNORs, we found two genes encoding for putative and almost identical GSNOR isoenzymes. One of the two, here named CrGSNOR1, was heterologously expressed and purified. The kinetic properties of CrGSNOR1 were determined and the high-resolution three-dimensional structures of the apo and NAD^+^-bound forms of the enzyme were solved. These analyses revealed that CrGSNOR1 has a strict specificity towards GSNO and NADH, and a conserved 3D-folding with respect to other plant GSNORs. The catalytic zinc ion, however, showed an unexpected variability of the coordination environment. Furthermore, we evaluated the catalytic response of CrGSNOR1 to thermal denaturation, thiol-modifying agents and oxidative modifications as well as the reactivity and position of accessible cysteines. Despite being a cysteine-rich protein, CrGSNOR1 contains only two solvent-exposed/reactive cysteines. Oxidizing and nitrosylating treatments have null or limited effects on CrGSNOR1 activity, highlighting a certain resistance of the algal enzyme to redox modifications. The molecular mechanisms and structural features underlying the response to thiol-based modifications are discussed.

**One-sentence summary:** GSNOR1 from *Chlamydomonas reinhardtii* displays an unusual variability of the catalytic zinc coordination environment and an unexpected resistance to thiol-based redox modifications

## INTRODUCTION

Nitric oxide (•NO) is a relatively stable free radical widely recognized as a signaling molecule in oxygenic phototrophs where it controls multiple physiological processes (*e.g*. development, stomatal closure, tolerance to metal toxicity, and adaptive response to abiotic and biotic stresses) (Neill et al., 2008; Bellin et al., 2013; Umbreen et al., 2018; Ageeva-Kieferle et al., 2019; Del Castello et al., 2019; Li et al., 2019; Kuo et al., 2020; Yu et al., 2020). The biological actions of •NO are mainly exerted by NO-derived reactive molecules through their ability to react with proteins and trigger the formation of post-translational modifications (PTMs) (Besson-Bard et al., 2008; Begara-Morales et al., 2016; Feng et al., 2019; Gupta et al., 2020). The major reaction consists in the reversible formation of a nitrosothiol (−SNO) between a NO moiety and a protein thiol (−SH), in a process named S-nitrosylation (Zaffagnini et al., 2019).

Protein S-nitrosylation has emerged as an important regulatory process in plants and hundreds of proteins have been identified as putative S-nitrosylated targets both *in vitro* and *in vivo* (Lindermayr et al., 2005; Astier et al., 2012; Zaffagnini et al., 2013; Morisse et al., 2014; Yu et al., 2014; Zaffagnini et al., 2014; Zaffagnini et al., 2016; Huang et al., 2019; Skelly et al., 2019). However, •NO itself cannot directly react with cysteine thiols, but can readily condense with oxygen leading to the formation of nitrogen dioxide (•NO_2_). Subsequently, •NO_2_ can react with •NO to form dinitrogen trioxide (N_2_O_3_) that can induce S-nitrosothiol formation by reacting with sulfur atoms of low-molecular weight thiols and protein cysteines (Zaffagnini et al., 2016). Considering the high intracellular concentration of reduced glutathione (GSH; γ-Glu-Cys-Gly) (1-5 mM; (Rouhier et al., 2008; Noctor et al., 2012)), nitrosoglutathione (GSNO) is suggested to be the most abundant intracellular low-molecular weight S-nitrosothiol (Airaki et al., 2011; Corpas et al., 2013). GSNO is a quite stable NO-carrying molecule that is considered as the major NO reservoir in both plant and animal cells (Broniowska et al., 2013; Corpas et al., 2013). In addition, GSNO can donate its NO moiety to protein cysteines through a trans-nitrosylation reaction (Zaffagnini et al., 2019). Due to GSNO reactivity, its intracellular concentration must be tightly regulated to avoid uncontrolled accumulation of S-nitrosylated proteins that might cause severe perturbations of cell metabolism and signaling. In animals, several enzymes were shown to catabolize GSNO, including thioredoxin (TRX) (Sengupta and Holmgren, 2012), glutaredoxin (GRX) (Ren et al., 2019), superoxide dismutase (SOD) (Okado-Matsumoto and Fridovich, 2007; Lushchak et al., 2009) nitrosoglutathione reductase (GSNOR) (Liu et al., 2001), human carbonyl reductase 1 (HsCBR1) (Bateman et al., 2008), and the recently described aldo-keto reductase family 1 member A1 (HsAKR1A1) (Stomberski et al., 2019). Unlike TRX, GRX, and SOD, which catalyze the reduction of GSNO yielding GSH and other NO-derived molecules as final products, GSNOR along with HsCBR1 and HsAKR1A1 catalyze the irreversible conversion of GSNO to N-hydroxysulfinamide (GSNHOH), an unstable intermediate that, in the presence of reduced glutathione (GSH), yields oxidized glutathione (GSSG) and hydroxylamine (Liu et al., 2001; Kubienova et al., 2013; Zaffagnini et al., 2016). For this reason, GSNOR acts as a scavenging system of intracellular GSNO, thereby indirectly influencing the extent of protein S-nitrosylation (Lindermayr, 2017; Jahnova et al., 2019). Consistently, yeast strains, mice, *Arabidopsis thaliana* and *Lotus japonicus* plants deficient for GSNOR exhibited increased levels of protein S-nitrosothiols (SNOs) (Liu et al., 2004; Feechan et al., 2005; Foster et al., 2009; Matamoros et al., 2020), while a decrease of SNO levels was observed in plants overexpressing GSNOR (Lin et al., 2012). Overall, these data suggest that GSNO positively correlates with S-nitrosylated proteins *in vivo*, and that GSNOR is an enzymatic scavenging system capable of regulating GSNO levels in different organisms including plants.

GSNOR belongs to the class-III alcohol dehydrogenase family and can be found in most bacteria and all eukaryotes including photosynthetic organisms (Liu et al., 2001). This enzyme was originally identified as a glutathione-dependent formaldehyde dehydrogenase and then reclassified as an S-(hydroxymethyl)glutathione (HMGSH) dehydrogenase. Lately, it was found to participate in GSNO catabolism by catalyzing GSNO reduction using NADH as electron donor (Jensen et al., 1998; Liu et al., 2001; Sakamoto et al., 2002; Kubienova et al., 2013). In photosynthetic organisms, GSNOR is generally localized in the cytoplasm and encoded by a single gene (Lee et al., 2008), with few exceptions including poplar, *Lotus japonicus* and *Chlamydomonas reinhardtii* which contain two GSNOR nuclear genes (Merchant et al., 2007; Xu et al., 2013; Cheng et al., 2015). Crystal structures show that GSNOR is a homodimeric protein containing two zinc ions per monomer having either a catalytic or a structural role (Sanghani et al., 2002; Kubienova et al., 2013; Jahnova et al., 2019).

Recently, several studies reported that Arabidopsis and poplar GSNOR undergo S-nitrosylation *in vivo* under conditions of increased endogenous NO levels (Frungillo et al., 2014; Cheng et al., 2015). Moreover, this modification affects GSNOR activity following exposure of Arabidopsis leaf extracts to NO-donors (Frungillo et al., 2014) and is controlled by GSH as proven by both *in vitro* and genetic studies *in vivo* (Zhang et al., 2020). The kinetics and structural effects of S-nitrosylation on GSNOR from Arabidopsis have been reported and the nitrosylated cysteine residues identified (Cys10, 271, and 370) (Guerra et al., 2016). The specific S-nitrosylation of Cys10 triggers AtGSNOR degradation through autophagy under hypoxic conditions (Zhan et al., 2018). In addition, the redox modification of Cys10 occurs through a trans-nitrosylation reaction involving catalase 3 (Chen et al., 2020). Differently, in the leguminosae *Lotus japonicus*, the two GSNOR isoforms were found to be target of S-nitrosylation without effect on protein catalysis (Matamoros et al., 2020). Plant GSNORs were also found to be inhibited by *in vitro* treatments with hydrogen peroxide (Kovacs et al., 2016; Ticha et al., 2017; Matamoros et al., 2020) or after exposure of Arabidopsis and *Baccaurea ramiflora* plants to the pro-oxidant herbicide paraquat and exogenous hydrogen peroxide, respectively (Bai et al., 2012; Kovacs et al., 2016). Altogether, these results suggest that most plant GSNORs are responsive to oxidative modifications and transient inhibition of their activity might represent an important mechanism to control GSNO accumulation with an ensuing impact on intracellular GSNO/SNO levels.

In green microalgae such as *Chlamydomonas reinhardtii*, NO signaling participates in the regulation of nutrients acquisition, photosynthetic efficiency, and other processes including autophagy and cell death (Sanz-Luque et al., 2013; Wei et al., 2014; Calatrava et al., 2017; Zalutskaya et al., 2018; De Mia et al., 2019; Kuo et al., 2020), making its understanding of particular interest for biotechnological purposes. Recently, GSNO reducing activity has been measured in *Chlamydomonas reinhardtii* extracts following exposure to salt stress (Chen et al., 2016), but the underlying enzymes along with their functional features are yet to be uncovered.

In this work, we identified the enzymatic systems catalyzing GSNO degradation in *C. reinhardtii* protein extracts. Genome mining confirmed the presence in Chlamydomonas of two nuclear-encoded genes for putative GSNOR isozymes sharing more than 99% of sequence identity. Algal *GSNOR1* (*Cre12.g543400*) was cloned and expressed, and its biochemical and structural features determined. Despite being rich in cysteine residues (16 Cys out of 378 total residues), CrGSNOR1 contains only two solvent-exposed/reactive cysteines and its activity is almost unaffected by *in vitro* oxidative and nitrosative treatments, suggesting that the algal enzyme is resistant to redox modifications. Based on our findings, we provide functional and structural insights into the response of CrGSNOR1 to cysteine-based modifications.

## MATERIAL AND METHODS

### Chemicals

Proteomics grade Trypsin Gold was obtained from Promega. Desalting columns (NAP-5 and PD-10) and N-[6-(Biotinamido)hexyl]-3’-(2’-pyridyldithio)proprionamide (HPDP-biotin) were purchased from GE Healthcare and Pierce, respectively. All chemicals were obtained from Sigma-Aldrich unless otherwise specified.

### Synthesis of S-nitrosoglutathione

GSNO was synthesized from commercial glutathione via an acid-catalyzed nitrosation reaction as previously described in (Hart, 1985). Briefly, commercial glutathione (3.065 g) was dissolved in 21 ml of a 476 mM hydrochloric acid solution and kept on ice. Sodium nitrite (0.691 g) was added at once and the mixture was kept under stirring for 45 min and protected from light. Then, 10 ml of acetone were added to the red slurry and kept under stirring for an additional 10 min. The slurry was filtered on a glass frit and the precipitate was washed with prechilled distilled water (4 x 20 ml), acetone (3 x 20 ml) and diethyl ether (3 x 20 ml). Water and solvent traces were removed under vacuum for 24 h and the powder (avg. yield 70%) was kept at −20°C in the presence of desiccant. GSNO purity was assessed by ^1^H-NMR and the concentration was determined spectrophotometrically using molar extinction coefficients of 920 M^−1^ cm^−1^ and 15.9 M^−1^ cm^−1^ at 335 nm and 545 nm, respectively.

### Cell culture, growth conditions and protein extraction

Conditions for Chlamydomonas cultures and protein extraction were adapted from (Morisse et al., 2014). Briefly, the Chlamydomonas D66 cell-wall-less strain (CC-4425 cw nit2-203mt+ strain) was grown in Tris-acetate phosphate (TAP) medium under continuous light (80 μE m^−2^ s^−1^) at 25 °C up to 4-5 x 10^6^ cells ml^−1^. Cultures were then pelleted (4000 *g*, 5 min) and resuspended in 50 mM Tris-HCl pH 7.9. Total soluble proteins were then extracted by three cycles of freeze/thaw in liquid nitrogen. The protein extract was then clarified by centrifugation (15000 *g* for 10 min at 4 °C) and protein concentration was assessed by BCA Protein Assay using bovine serum albumin (BSA) as standard.

### NAD(P)H-dependent GSNO reductase activity in protein extracts

The NAD(P)H-dependent GSNO reductase activity was measured adding variable amounts of freshly prepared protein extracts (0.125-1 mg) in a reaction mixture (1 ml) containing 50 mM Tris-HCl pH 7.9, 0.2 mM NAD(P)H and 0.4 mM GSNO. The activity was determined spectrophotometrically following NAD(P)H oxidation at 340 nm using a molar extinction coefficient of 7060 M^−1^ cm^−1^ at 340 nm, which includes both NAD(P)H and GSNO absorbance. The linear rate of the reaction was corrected with a reference rate without GSNO. Activity measurements were performed at least in three biological triplicates using 1 cm path length cuvettes.

### Thiol-modifying treatments and thermal stability of protein extracts

Freshly prepared protein extracts (500 μg) were incubated at 25 °C in 50 mM Tris-HCl, pH 7.9 in the presence of 1 mM N-ethylmaleimide (NEM) or 1 mM and methyl methanethiosulfonate (MMTS). At the indicated times, aliquots (10-50 μl) were withdrawn to carry out activity measurements as described above. Control experiments were performed by incubating protein extracts in the presence of 2 mM reduced DTT. Thermal stability was carried out by incubating protein extracts (500 μg) for 5 min from 40 °C up to 80 °C with 10 °C increments. Subsequently, protein samples were centrifuged (15000 *g* for 5 min at 4 °C) to remove precipitated proteins, and the NAD(P)H-dependent activities were monitored as described above. Control experiments were performed by incubating protein extracts at 25 °C following the centrifugation step.

### Cloning, expression and purification of CrGSNOR1

The coding sequence for *CrGSNOR1* (locus *Cre12.g543400*) was amplified by standard RT-PCR on Chlamydomonas total RNA extracts using a forward primer introducing an *Nde*I restriction site (in bold) at the start codon: 5’-CATGCC**CATATG**TCGGAAACTGCAGGCAAG-3’ and a reverse primer introducing a *BamHI* restriction site (in bold) downstream of the stop codon: 5’-CATGCC**GGATCC**CTAGAACGTCAGCACACA-3’. *CrGSNOR1* was cloned in a modified pET-3c vector (Pasquini et al., 2017) containing additional codons upstream of the *NdeI* site to express a His-tagged protein with seven N-terminal histidines. The sequence was checked by sequencing. Recombinant CrGSNOR1 was produced using the pET-3c-His/BL21 expression system. Bacteria were grown in LB medium supplemented with 100 μg ml^−1^ ampicillin at 37 °C and the production was induced with 100 μM isopropyl β-D-1-thiogalactopyranoside overnight at 30°C. Cells were then harvested by centrifugation (5000 *g* for 10 min) and resuspended in 50 mM Tris-HCl pH 7.9. Cell lysis was performed using a French press (6.9 x 10^7^ Pa) and cell debris were removed by centrifugation (5000 *g* for 15 min). To avoid nucleic acids contamination, the sample was incubated with RNase (0.01 mg ml^−1^) and DNAse (0.04 U ml^−1^) for 30 min at RT under mild shaking. The supernatant was then centrifuged at 15000 *g* for 30 min and applied onto a Ni^2+^ Hitrap chelating resin (HIS-Select Nickel Affinity Gel; Sigma-Aldrich) equilibrated with 30 mM Tris-HCl pH 7.9 containing 500 mM NaCl (TN buffer) and 5 mM imidazole. The recombinant protein was purified according to the manufacturer’s instructions. The molecular mass and purity of the protein were analyzed by SDS-PAGE after desalting with PD-10 columns equilibrated with 30 mM Tris-HCl pH 7.9. The concentration of CrGSNOR1 was determined spectrophotometrically using a molar extinction coefficient at 280 nm (ε_280_) of 40910 M^−1^ cm^−1^. The resulting homogeneous protein solutions were stored at −20 °C.

### Enzymatic assays for GSNOR activities

The catalytic activity of purified CrGSNOR1 was measured spectrophotometrically as described above. The reaction was initiated by the addition of CrGSNOR1 at a final concentration ranging from 5 to 50 nM. The NADH-dependent activity of CrGSNOR1 was also assayed in the presence of oxidized glutathione (0.4 or 4 mM) or 0.2 mM NADPH instead of GSNO or NADH, respectively. S-(hydroxymethyl)glutathione (HMGSH) oxidation by CrGSNOR1 was assessed following the procedure described in (Sanghani et al., 2006). Briefly, the activity was determined spectrophotometrically following NAD^+^ reduction in a reaction mixture containing 50 mM Tris-HCl pH 7.9, 0.2 mM NAD^+^ and 1 mM HMGSH. The activity was measured as the increase in absorbance at 340 nm using a ε_340_ of 6220 M^−1^ cm^−1^.

### Kinetic properties of CrGSNOR1

Steady-state kinetic analysis was accomplished by varying the concentrations of NADH (0.005-0.2 mM) at a fixed GSNO concentration (0.4 mM) and the concentration of GSNO (0.0125-0.4 mM) at a fixed concentration of NADH (0.2 mM). The reaction was started by adding 25 nM CrGSNOR1. Three independent experiments were performed at each substrate concentration and apparent kinetic parameters (*K*’_m_ and *k*’_cat_) were calculated by nonlinear regression using the Michaelis-Menten equation with the program CoStat (CoHort Software, Monterey, CA).

### Thermal stability of purified CrGSNOR1

The thermostability of purified CrGSNOR1 (5 μM) was assessed by measuring protein activity after 30 min incubation of the enzyme at temperatures ranging from 25 °C up to 75 °C with 5 °C increments. Kinetics of CrGSNOR1 aggregation were assessed by measuring the increase of turbidity at 405 nm. CrGSNOR1 samples were incubated in 30 mM Tris-HCl, pH 7.9 at the indicated temperatures in a low-protein-binding 96-well plate. Samples were monitored at interval times and turbidity was measured using a plate reader (Victor3 Multilabeling Counter; Perkin-Elmer).

### Thiol-modifying treatments of CrGSNOR1

Treatments were performed at room temperature by incubating purified CrGSNOR1 (5 μM) in 50 mM Tris-HCl, pH 7.9 in the presence of NEM and MMTS at 1 mM. After 30 min incubation, aliquots were withdrawn to assay GSNOR activity as described above.

### Alkylation of CrGSNOR1 by maleimide-based reagents

Recombinant CrGSNOR1 (10 μM) was incubated in 30 mM Tris-HCl, pH 7.9 at room temperature in the presence of either 1 mM N-ethyl maleimide (100 mM stock solution prepared in water) or 1 mM Biotin-maleimide (50 mM stock solution prepared in DMSO). At indicated time points (20, 30, 60, 90 min) DTT (10 mM) was added to quench maleimide derivatives.

### In-solution trypsin digestion

Alkylated CrGSNOR1 (100 μl) was immediately desalted by gel filtration using NAP-5 columns equilibrated in water as recommended by the supplier. Then, the desalted protein samples (ca. 500 μl) were concentrated using a SpeedVac concentrator. CrGSNOR1 concentration was determined spectrophotometrically before a 3 h digestion step with trypsin (1:20 (w/w) enzyme:substrate ratio) in 25 mM ammonium bicarbonate (AMBIC). Trypsin digestion was stopped either by heating at 95 °C for 3 min or by ultrafiltration using 0.5 ml Amicon Ultra centrifugal devices (20 kDa MWCO, Millipore). A five microliters aliquot was kept for MALDI-TOF MS analysis and the rest was used for the enrichment of biotinylated peptides by affinity chromatography.

### Affinity purification of cysteinyl peptides alkylated by Biotin-maleimide

Affinity purification was performed as previously described in (Pérez-Pérez et al., 2017) with slight modifications. Briefly, around 75 μl of monomeric avidin agarose (Pierce) were packed into a gel-loading tip and further equilibrated with 200 mM NaCl in 25 mM AMBIC (loading buffer). Peptide mixture was supplemented with 200 mM NaCl before loading by centrifugation (20 °C, 1 min, 40 g). The flow through was kept and reloaded three times. Then, avidin agarose was extensively washed by centrifugation with 4 x 150 μl of loading buffer and 4 x 150 μl of 25 mM AMBIC in 20% methanol. Peptides retained onto the packed monomeric avidin column were eluted using 150 μl of 0.4% trifluoroacetic acid (TFA) in 30% acetonitrile (ACN) and were directly analyzed by MALDI-TOF without further treatment.

### MALDI-TOF MS analyses

Mass spectrometry experiments were performed as previously described in (Marchand et al., 2019; Shao et al., 2019). Briefly, for analysis of intact proteins by mass spectrometry, 1 μl of protein sample (previously quenched with DTT as described above) was taken and mixed with 2 μl of a saturated solution of sinapinic acid in 30/0.3 ACN/TFA. Two microliters of this premix were spotted onto the sample plate and allowed to dry under a gentle air stream at room temperature. Spectra were acquired in positive linear mode on an Axima Performance MALDI-TOF/TOF mass spectrometer (Shimadzu-Kratos, Manchester, UK) with a pulse extraction fixed at 50000. Mass determination was performed after external calibration using mono-charged and dimer ions of yeast enolase.

### Treatments of CrGSNOR1 with oxidizing or nitrosylating agents

Oxidizing and nitrosylating treatments were performed at 25 °C by incubating purified CrGSNOR1 (5 μM) in 50 mM Tris-HCl, pH 7.9 in the presence of 1 mM hydrogen peroxide (H_2_O_2_), or 1 mM diamide (TMAD), or 2 mM GSNO, or SNAP (0.2 and 2 mM). After 30 min incubation, an aliquot was withdrawn, and enzyme activity was assayed as described above. Reactivation of SNAP-treated CrGSNOR1 was carried out after 20 min incubation in the presence of 10 mM DTT.

### Biotin Switch Technique

Purified CrGSNOR1 was incubated in TEN buffer (30 mM Tris-HCl pH 7.9, EDTA 1 mM, NaCl 100 mM) in the presence of 2 mM GSNO or 2 mM SNAP for 30 minutes in the dark at 25 °C. The extent of protein nitrosylation was assessed by following the procedure described in (Zaffagnini et al., 2013). After nitrosylation treatments, proteins (~1 mg ml^−1^) were precipitated with two volumes of 80% cold acetone at −20 °C during 20 min and pelleted by centrifugation at 4 °C for 10 min at 15,000 *g*. The pellet was resuspended in TENS buffer (30 mM Tris-HCl pH 7.9, 1 mM EDTA, 100 mM NaCl and 1% SDS) supplemented with a cocktail of alkylating reagents (10 mM iodoacetamide, 10 mM N-ethylmaleimide), to allow blocking of free thiols. After 30 min incubation at 25 °C under shaking, the samples were acetone precipitated, as described above, to remove unreacted alkylating reagents. After resuspension in TENS buffer, proteins were incubated in the presence of 40 mM ascorbate and 1 mM N-[6-(Biotinamido)hexyl]-3’-(2’-pyridyldithio)propionamide (HPDP-biotin) for 30 min. This step allows reduction of S-nitrosylated cysteines and their derivatization with biotin. Proteins were then acetone precipitated to remove unreacted labelling compounds, pelleted by centrifugation as above and resuspended in TENS buffer. All steps were performed in the dark. After the final precipitation, proteins were quantified using the bicinchoninic acid assay, separated by non-reducing SDS-PAGE and transferred onto nitrocellulose membranes. Protein loading and transfer were assessed by Ponceau staining of the membrane. Proteins were then analyzed by western blotting using a primary anti-biotin antibody (1:5,000 dilution; Sigma-Aldrich) and an anti-mouse secondary antibody coupled to peroxidase (1:10,000 dilution; Sigma-Aldrich). Signals were visualized by enhanced chemiluminescence as described previously (Zaffagnini et al., 2012). All BST assays included a negative control where ascorbate was omitted to prevent reduction of S-nitrosothiols and subsequent biotinylation.

### Quaternary structure determination

Gel filtration analysis was performed on a Superdex 200 HR10/300 GL column (GE Healthcare) connected to an ÅKTA Purifier system (GE Healthcare), previously calibrated with standard proteins, namely ferritin (440 kDa), aldolase (158 kDa), ovalbumin (43 kDa), and chymotrypsinogen A (25 kDa), as described in (Pasquini et al., 2017). The column was equilibrated with 50 mM Tris-HCl, pH 7.5 and 150 mM KCl. The loading volume of CrGSNOR1 samples was 0.25 ml at a concentration above 1 mg ml^−1^ and fractions of 0.5 ml were collected at a flow rate of 0.5 ml min^−1^. DLS measurements were performed employing a Malvern Nano ZS instrument equipped with a 633 nm laser diode (Zaffagnini et al., 2019). Samples consisting of CrGSNOR1 (5-50 μM) in 30 mM Tris-HCl, pH 7.9 were introduced in disposable polystyrene cuvettes (100 μl) of 1 cm optical path length. The width of DLS hydrodynamic radius distribution is indicated by the polydispersion index. In the case of a monomodal distribution (Gaussian) calculated by means of cumulant analysis, PdI = (σ/Z_avg_)^2^, where σ is the width of the distribution and Z_avg_ is the average radius of the protein population. The reported hydrodynamic radii (R_h_) have been averaged from the values obtained from five measurements, each one being composed of ten runs of 10 seconds.

### Crystallization and Data Collection

The apo- and holo-forms of CrGSNOR1 were crystallized using the hanging drop vapor diffusion method at 20 °C. The drop was obtained by mixing 2 μl of 5 mg ml^−1^ protein solution in 30 mM Tris-HCl, pH 7.9, 1 mM EDTA and only for the holo-enzyme 1 mM NAD^+^, and an equal volume of a reservoir solution containing 0.1 M Tris-HCl pH 8.5, 0.1 M MgCl_2_ or Mg(CH_3_CO_2_)_2_, and 12-15% w/v PEG 20K or 12% w/v PEG 8K as precipitant. Crystals with a rod-like morphology appeared after about 10 days. The crystals were fished, briefly soaked in a cryo-solution containing the reservoir components plus 20% v/v PEG 400, and then frozen in liquid nitrogen. Diffraction data were collected at 100 K using the synchrotron radiation of the beamline ID23-1 at ESRF (Grenoble, France) for apo-CrGSNOR1 and of the XRD1 beamline at Elettra (Trieste, Italy) for NAD^+^-CrGSNOR1. Data collections were performed with a wavelength of 1.0 Å for both crystals, an oscillation angle (Δϕ) of 0.1° and a sample-to-detector distance (d) of 385.62 mm (Pilatus 6M) for the apo-enzyme, while Δϕ=0.3° and d = 260.00 mm (Pilatus 2M) for the NAD^+^-enzyme. The images were indexed with XDS (Kabsch, 2010) and scaled with AIMLESS (Evans and Murshudov, 2013) from the CCP4 package. The unit cell parameters and the data collection statistics are reported in Supplemental Table 1.

### Structure Solution and Refinement

Apo-CrGSNOR1 structure was solved by molecular replacement with the program MOLREP (Vagin and Teplyakov, 2010) using the coordinates of apo-GSNOR from tomato as search model (PDB code 4DLA; (Kubienova et al., 2013)). Three dimers were placed in the asymmetric unit consistently with the calculated Matthews coefficient (Matthews, 1968) equal to 2.4 Å^3^ Da^−1^ for six molecules in the asymmetric unit and corresponding to a solvent content of 48%. The refinement was performed with REFMAC 5.8.0135 (Murshudov et al., 2011) selecting 5% of reflections for R_free_, and the manual rebuilding with Coot (Emsley and Cowtan, 2004). Water molecules were automatically added and, after a visual inspection, confirmed in the model only if contoured at 1.0 σ on the (2*F*_o_ – *F*_c_) electron density map and they fell into an appropriate hydrogen-bonding environment. Several PEG molecules, chloride and magnesium ions coming from the crystallization solution were identified and added to the model. The last refinement cycle was performed with PHENIX (Adams et al., 2010).

Since the NAD^+^-CrGSNOR1 crystal was isomorphous with the apo-form, the final coordinates of apo-CrGSNOR1 were directly used for refinement providing R and R_free_ values of 0.23 and 0.28, respectively. The calculated 2*F*_o_ – *F*_c_ and *F*_o_ – *F*_c_ electron density maps revealed a clear density for NAD^+^ in each monomer that was added to the structural model. The refinement of the holo-structure was performed as described for the apo-form. Refinement statistics are reported in Supplemental Table 1. The stereo-chemical quality of the models was checked with Molprobity (Chen et al., 2010). Molecular graphics images were generated using PyMOL (The PyMOL Molecular Graphics System, Schrödinger, LLC) and Ligplot (Wallace et al., 1995).

### Secondary structure analysis

The secondary structure of apo-CrGSNOR1 was investigated by means of circular dichroism (CD) spectroscopy. Samples of apo-CrGSNOR1 (10.7 μM) were prepared in 30 mM Tris-HCl, pH 7.9 and quantified by spectrophotometric analysis at 280 nm in a 1 cm cell (Pace et al., 1995). Oxidized apo-CrGNSOR1 was obtained by treatment with 1 mM H_2_O_2_. Far-UV CD spectra (260–190 nm) were measured at room temperature on a J-810 spectropolarimeter (Jasco, Japan), using a QS-quartz cylindrical cell with 0.2 mm optical pathlength (Hellma Analytics, Germany), a 1 nm spectral bandwidth, a 20 nm/min scanning speed, a 4 s data integration time, a 0.2 nm data interval and an accumulation cycle of 3 scans. The resulting CD spectra were blank-corrected and converted to molar units per residue (Δεres, in M^−1^ cm^−1^). The estimation of the secondary structure from the CD spectra of apo-CrGSNOR1 was performed using the CONTIN-LL algorithm (van Stokkum et al., 1990) and the 48-protein reference set 7 (Sreerama and Woody, 2000) available on the DichroWeb web server (http://dichroweb.cryst.bbk.ac.uk/) (Whitmore and Wallace, 2004).

### Accession numbers

Atomic coordinates and structure factors have been deposited in the Protein Data Bank (www.wwpdb.org) under PDB ID codes XXXX and XXXX for apo and NAD^+^-CrGSNOR1, respectively.

## RESULTS

### Distinct NADPH- and NADH-dependent enzymatic systems catalyze GSNO reduction in *C. reinhardtii*

To determine whether *C. reinhardtii* contains enzymatic systems able to catabolize GSNO, we examined GSNO reduction in the presence of NADPH or NADH by monitoring the decrease in absorbance at 340 nm. Chlamydomonas protein extracts were found to catalyze GSNO reduction using both cofactors and the relative activities correlated with protein content (Figure 1A and 1B). The NADPH-dependent specific activity (75.9 ± 10.0 nmol min^−1^ mg^−1^) was around two-fold higher compared to that measured in the presence of NADH (32.9 ± 2.3 nmol min^−1^ mg^−1^). To investigate whether the NADPH- and NADH-dependent activities are due to different enzymatic systems, we sought to find conditions that allowed uncoupling them. We first compared the thermal stability of the two enzymatic activities as it is well established that enzymes can exhibit very different sensitivity to temperature (Bischof and He, 2005). After incubation of protein extracts at varying temperatures ranging from 25 °C to 80 °C, we measured GSNO degradation in the presence of both cofactors. The NADPH-dependent activity was resistant to temperature up to 70 °C and strong inactivation was only achieved at 80 °C (Figure 1C). By contrast, NADH-dependent GSNO degradation exhibited a much higher sensitivity to heating, retaining 85%, 20% and 5% residual activity at 50 °C, 60 °C and 70 °C, respectively (Figure 1D).

**Figure 1.**
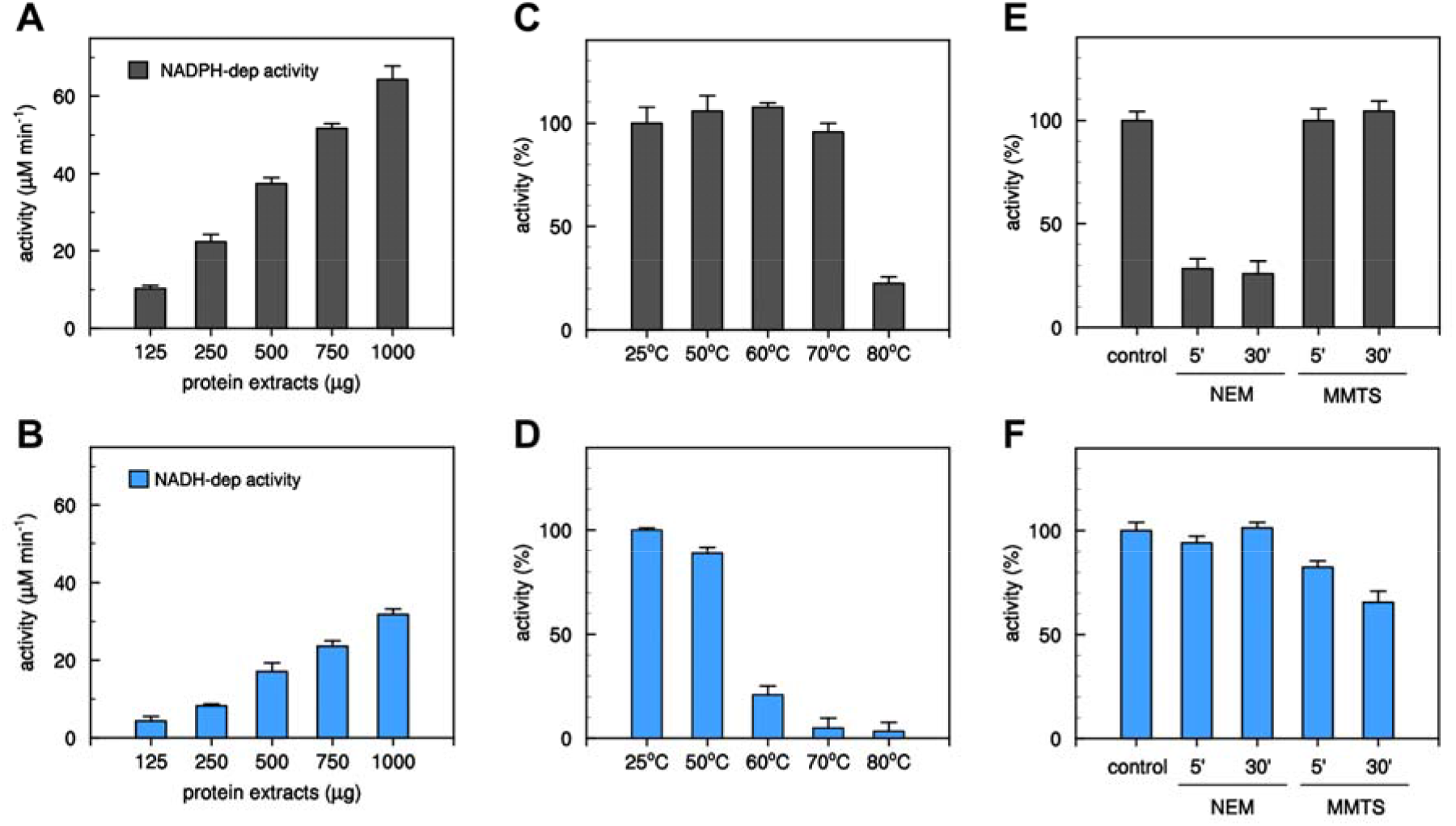
Measurements of GSNO-reducing activities from Chlamydomonas cell extract. (**A**) and (**B**) Determination of GSNO-reducing activity by variable amounts of protein extract from *Chlamydomonas* cell culture in the presence of NAD(P)H (NADPH, black bars; NADH, white bars). Data represented the mean ± SD calculated from three biological replicates (*n* = 3). (**C**) and (**D**) Thermal sensitivity of NAD(P)H-dependent GSNO reducing activity. Protein extracts (500 μg) were exposed to various temperatures and after incubation the NAD(P)H-dependent activities were assayed (NADPH, black bars; NADH, white bars). (**E**) and (**F**) Alkylation sensitivity of NAD(P)H-dependent GSNO reducing activity. Protein extracts were exposed for 5 or 30 min to 1 mM alkylating agents (NEM or MMTS) and after incubation the NAD(P)H-dependent activity was assayed (NADPH, black bars; NADH, white bars). For panels **C**-**F**, values are expressed as percentage of activity measured under control conditions (see Material and Methods) and are represented as mean percentage ± SD of three biological replicates (*n* = 3).

Further analyses were conducted aimed at investigating the response of NAD(P)H-dependent GSNO degrading activities to thiol-modifying agents such as N-ethylmaleimide (NEM) and methyl methanethiosulfonate (MMTS). These two compounds share a strong reactivity towards cysteine residues, but while NEM induces irreversible alkylation, MMTS reacts with sulfhydryl groups (−SH) forming a mixed disulfide (-S-S-CH_3_, dithiomethane). In addition, NEM exclusively reacts with accessible cysteine residues while MMTS can also react with metal-coordinating cysteine thiols (D’Ordine et al., 2012). The exposure of protein extracts to NEM led to a strong and rapid inactivation of the NADPH-dependent activity whereas no effect was observed when we assayed GSNO reduction in the presence of NADH (Figure 1E and 1F). By contrast, MMTS had no significant effect on the NADPH-dependent activity whereas it induced a partial decrease of NADH-dependent activity, retaining ~60% residual activity after 30 min incubation (Figure 1E and 1F).

Based on these findings, we can sustain that Chlamydomonas protein extracts contain at least two distinct GSNO-reducing enzymatic systems exhibiting specific cofactor dependence and different sensitivities to thermal denaturation and cysteine-modifying molecules.

### The Chlamydomonas genome contains two genes encoding nearly identical GSNOR isoforms

Since plant and non-plant GSNORs are known to specifically use NADH as electron donor, we sought to establish that the algal enzymatic system catalyzing NADH-dependent GSNO degradation could be ascribed to a GSNOR ortholog. Blast searches using GSNOR sequences from diverse sources revealed the presence of two *GSNOR* genes in the Chlamydomonas nuclear genome (v5.5). The two genes were annotated as formaldehyde dehydrogenases and we name them here *GSNOR1* (*Cre12.g543400*) and *GSNOR2* (*Cre12.g543350*). The two genes are most probably the result of a recent duplication, as they are adjacent and code for almost identical proteins (~99% sequence identity, Supplemental Figure 1). Multiple sequence alignments revealed that Chlamydomonas GSNORs (CrGSNORs) show 70% and 65% sequence identity with structurally solved GSNORs from land plants (*i.e. Arabidopsis thaliana* and *Solanum lycopersicum*) and human cells, respectively (Figure 2). Comparison of CrGSNOR sequences with GSNORs from different plant and non-plant species showed a similar amino acid conservation ranging from 54% to 72% sequence identity apart from GSNOR from the green alga *Volvox carteri* (90% identity) (Supplemental Figure 2). The residues involved in the coordination of both catalytic and structural zinc ions are fully conserved, and this also applies to residues participating in the stabilization of the cofactor NAD(H) (Figure 2). Based on the high sequence identity among analyzed GSNORs, we can hypothesize that algal GSNORs represent the enzymes responsible for the NADH-dependent GSNO reduction detected in Chlamydomonas protein extracts. To confirm this hypothesis, we investigated the structural and functional properties of CrGSNORs by focusing our attention on isoform 1 (CrGSNOR1).

**Figure 2.**
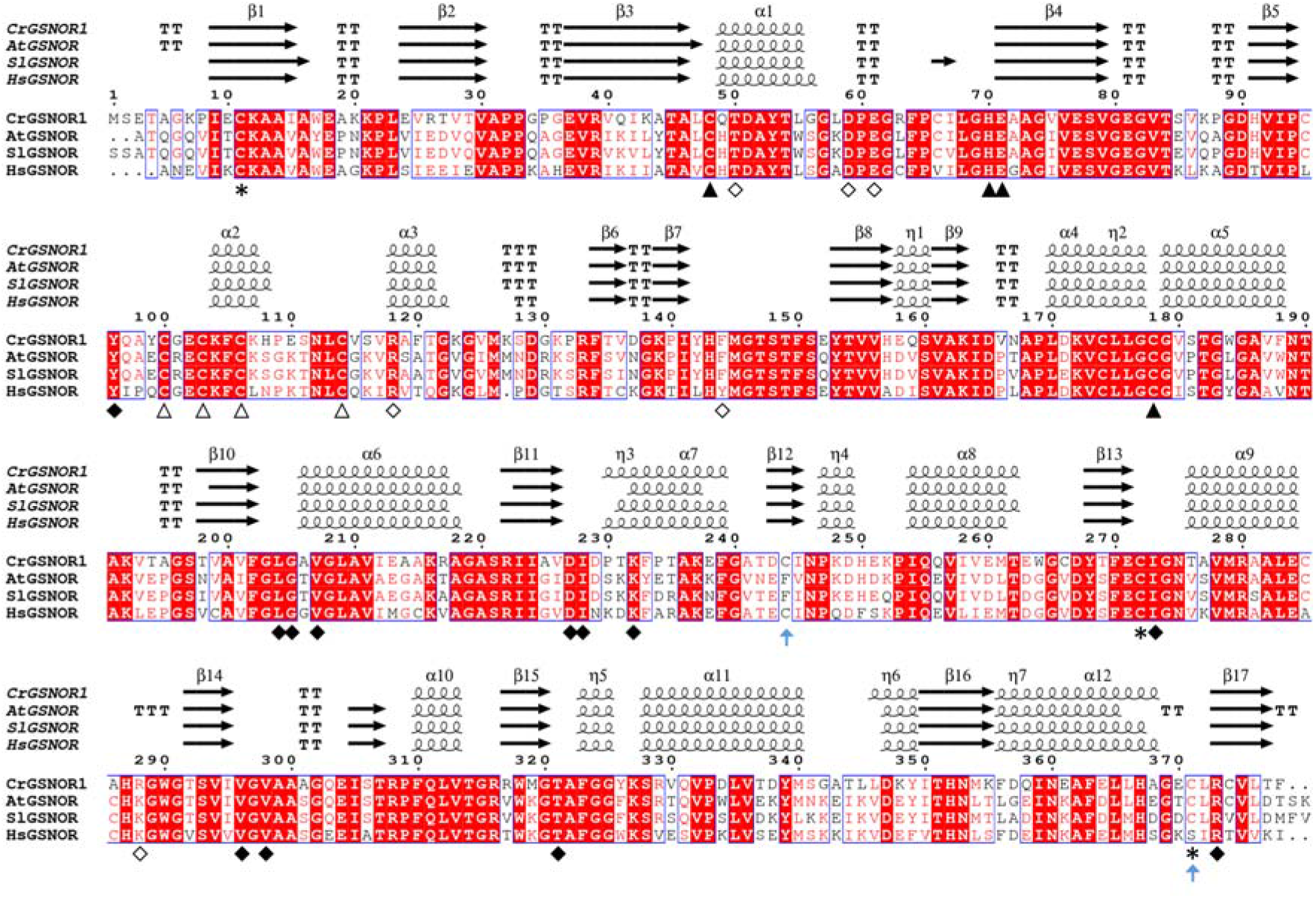
Primary and secondary structure alignment of 3D-solved GSNORs. The alignment was performed with Espript (http://espript.ibcp.fr) (Robert and Gouet, 2014) using the sequence and the structure of CrGSNOR1 (this work); GSNOR from *Arabidopsis thaliana* (AtGSNOR, PDB ID 3UKO); GSNOR from *Solanum lycopersicum* (SlGSNOR, PDB ID 4DLB), GSNOR from *Homo sapiens* (HsGSNOR, PDB ID 1M6H). The conserved residues are shown in red background; blue boxes represent conserved amino acid stretches (>70%). Residues with similar physico-chemical properties are indicated in red. α-helices, β-strands and 3_10_-helices are marked with α, β, η respectively. β-turns and α-turns are represented by TT and TTT, respectively. Residues coordinating the catalytic and structural zinc atom are indicated by closed and open triangles, respectively. Closed and open diamonds denote residues interacting with the cofactor and substrate, respectively. An asterisk indicates putative cysteine targets of S-nitrosylation in AtGSNOR while a light-blue arrow indicates accessible cysteine residues in CrGSNOR1. The primary sequence alignment was made using Clustal Omega (Sievers et al., 2011).

### CrGSNOR1 is a homodimeric protein displaying a conserved folding

To gain insight into the structural features of CrGSNOR1, we heterologously expressed the enzyme in *E. coli* as a 386 amino acids polypeptide (full-length protein plus the MHHHHHHH peptide at the N-terminus) and purified it to homogeneity by Ni^2+^ affinity chromatography. The purified protein migrated as a single band of ~40 kDa on SDS-PAGE under both reducing and non-reducing conditions (Supplemental Figure 3A), and MALDI-TOF mass spectrometry confirmed that recombinant CrGSNOR1 had the expected molecular mass of 41500.6 Da (Supplemental Figure 3B). Gel filtration and DLS analyses were conducted to determine the oligomerization state of CrGSNOR1. The enzyme eluted as a single symmetric peak with an apparent molecular mass of 96.4 ± 6.1 kDa and the elution profile at 280 nm perfectly correlated with GSNOR activity (Supplemental Figure 3C). These results clearly indicate that CrGSNOR1 protein is a non-covalent homodimer as further confirmed by DLS analysis that reported a hydrodynamic radius of 4.14 ± 0.2 nm, corresponding to an apparent molecular mass of 93.6 ± 4.3 kDa.

The dimeric fold was chiefly established by solving the crystal structure of CrGSNOR1 under both apo- and holo-form (NAD^+^-CrGSNOR1) at a resolution of 1.8 and 2.3 Å, respectively (Figure 3A and 3B). The apo- and holo-enzymes showed an identical crystalline packing with three dimers in the asymmetric unit and a similar overall structure with root mean square deviation (rmsd) values ranging from 0.20 to 0.86 Å and from 0.33 to 0.97 Å for monomers and dimers superimposition, respectively. Since similar rmsd values were obtained in the superimposition among the six monomers or three dimers of the same apo- or holo-form, we can conclude that the observed differences are mainly related to a conformational intrinsic variability of CrGSNOR1 molecules rather than to specific conformational changes between apo- and holo-structure. The comparison of CrGSNOR1 with other structurally known GSNORs (*i.e*. human, tomato and Arabidopsis GSNORs) clearly indicates a folding conservation with an almost identical secondary structure composition (Figure 2) (Sanghani et al., 2006; Kubienova et al., 2013; Xu et al., 2013). The mean rmsd values for dimers superimposition of apo-CrGSNOR1 with tomato apo-enzyme (PDB code 4DLA) is 0.83 Å and similar values (0.92 and 1.03 Å) were obtained when NAD^+^-CrGSNOR1 was superimposed to holo-enzymes from tomato (PDB code 4DL9) and Arabidopsis (PDB code 4JJI), respectively. The comparison with human (Hs) apo- and holo-CrGSNOR gave rmsd values within the same range (0.92 and 0.84 Å, respectively). All GSNOR structures known so far are thus very similar, and the differences between species are, in terms of rmsd, comparable to the differences among CrGSNOR1 dimers of the same asymmetric unit.

**Figure 3.**
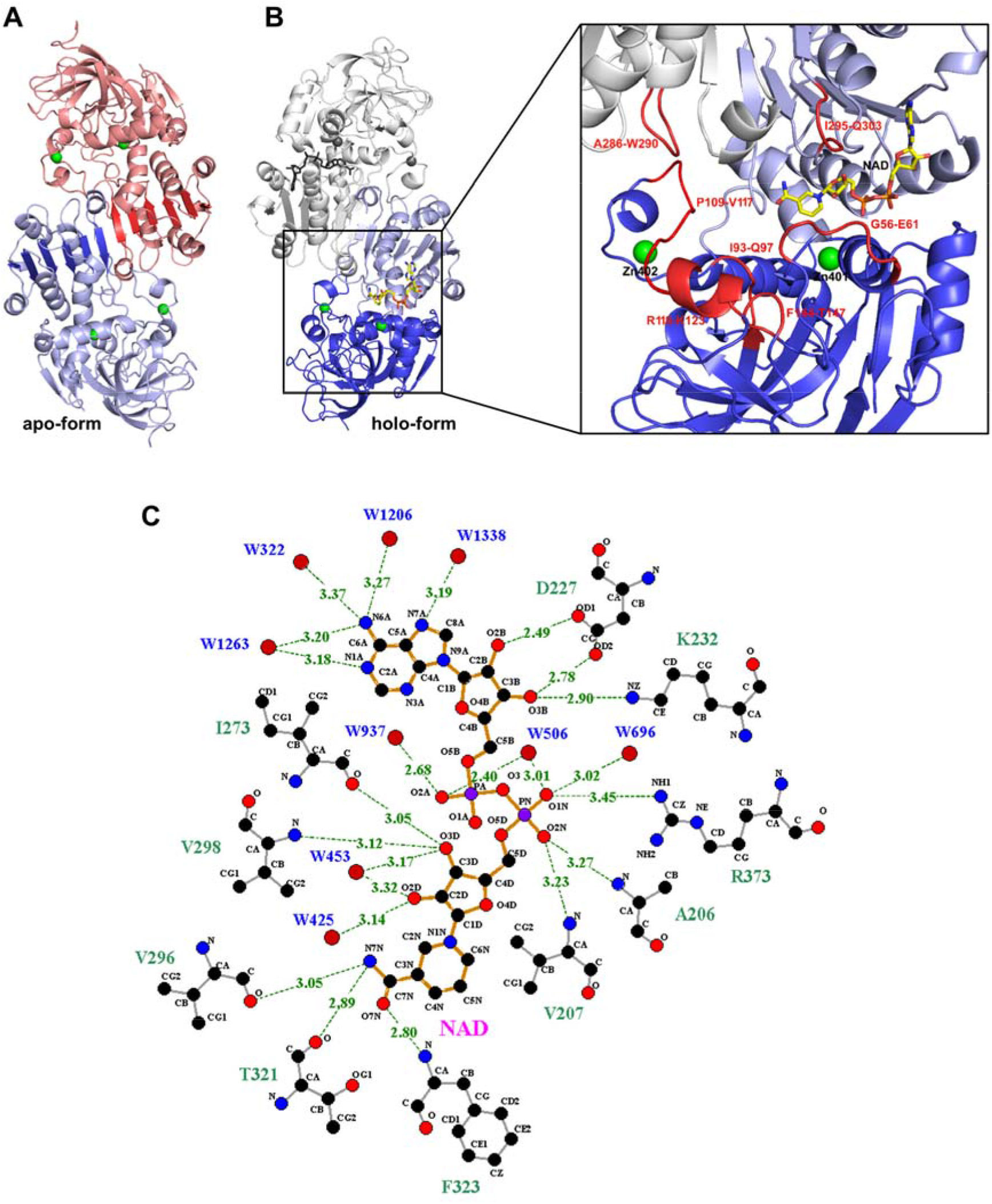
Crystal structure of apo- and NAD^+^-CrGSNOR1. (**A**) Overall folding of dimeric apo-CrGSNOR1. The two subunits are shown in salmon and light blue and the zinc ions of each subunit as green spheres. The six β-strand of each subunit forming a continuous β-sheet at the dimer interface are highlighted in red and blue. **(B)** Overall folding of dimeric NAD^+^-CrGSNOR1. NAD^+^ shown in sticks occupies the cofactor-binding domain (in light blue; residues 178-326) characterized by the typical Rossman fold. The larger catalytic domain (in blue; residues 1-177 and 327-377) comprises the metal ion sites. The active site of CrGSNOR1 (zoom) is located between the catalytic and cofactor-binding domains and is formed by the loops and the α-helix highlighted in red. (**C**) Hydrogen bond and salt-bridge interactions (up to 3.5 Å) of NAD^+^ with protein residues and water molecules.

The structural homology of CrGSNOR1 with other known GSNORs also embraces subunit composition. Indeed, each subunit is composed of a large catalytic domain comprising residues 1-177 and 327-377, and a smaller cofactor-binding domain (residues 178-326, Figure 3B). The latter domain shows the typical Rossman fold formed by a six-stranded parallel β-sheet sandwiched among six α-helices and an additional β-strand. This domain forms the internal dimer interface and is oriented in such a way that the six-stranded β-sheets of each subunit form a continuous β-sheet (Figure 3A). The cofactor is stabilized by several hydrogen bonds with protein residues and water molecules, and a unique electrostatic interaction established between its nicotinamide phosphate group and Arg373 (Figure 3C). The adenine ring is sandwiched between two isoleucine residues (Ile228 and Ile272) but does not form short interactions (< 3.5 Å) with protein residues (Figure 3C and Supplemental Figure 4A). The nicotinamide ring is kept in place by hydrophobic interactions with two valines (Val207 and Val298) and the methyl group of Thr182, and hydrogen bonds between its terminal amide group and the backbone carbonyl group of Val296 and Thr321, and amino group of Phe323 (Figure 3C and Supplemental Figure 4A).

### The catalytic domain of CrGSNOR1 allocates both the catalytic and structural zinc ions

The catalytic domain of CrGSNOR1 contains two zinc ions. One zinc ion (Zn402) is thought to have a structural role and it is coordinated with a tetrahedral geometry by four cysteine residues (Cys100, 103, 106 and 114) in both apo- and holo-forms (Figure 4A). The second zinc ion (Zn401) lies in the active site and has a catalytic role as a Lewis acid, activating the functional group of the substrate. In NAD^+^-CrGSNOR1, it is coordinated with a tetrahedral geometry involving Cys48, Cys178, His70, and Glu71 (Figure 4B). The identical geometry is maintained in one out of six subunits of the apo-structure (subunit F) with Glu71 replaced by a water molecule (or a hydroxide ion) (Figure 4C). In the other subunits, the metal ion is coordinated by five ligands comprising the four aforementioned residues and one water molecule in chains A, B, D and E (Figure 4D) or Cys48, Cys178, His70 and two water molecules in chain C (Figure 4E). This penta-coordination formed a distorted trigonal bipyramidal geometry with the oxygen ligands from Glu71 and/or water molecules in the axial positions (*i.e*. perpendicular to the equatorial plane), while the nitrogen from His70 and the two sulfur ligands from Cys48 and Cys178 are found on the equatorial plane forming 120° angles. In both subunits F and C, the metal center lies at more than 4 Å from Glu71 having its carboxylic group electrostatically interacting with Arg373 (Figure 4C and 4E). This Glu-Arg salt-bridge is conserved also in the subunits where Glu71 participates in Zn^2+^ coordination (Figure 4B and 4D). When the cofactor binds to the enzyme, Arg373 slightly moves toward the cofactor phosphate groups weakening the interaction with Glu71 that preferentially coordinates the zinc ion (Figure 4B). However, in two subunits of the NAD^+^-structure (C and F subunits) the distance Glu71-Zn401 is between 3.5-3.8 Å. Subunits superimposition shows that the increased Glu71-Zn401 distance observed in C and F subunits of both apo- and holo-forms is due to a 2-3 Å displacement of the zinc ion away from the glutamate toward the substrate-binding site (Supplemental Figure 5A and 5B) in a position superimposable to the catalytic zinc ion in HsGSNOR complexed with NADH and S-(hydroxymethyl)glutathione (HMGSH) (Sanghani et al., 2002) (Supplemental Figure 5C). The reversible association of the catalytic zinc ion to Glu71 (*i.e*. far in apo-structure, close in holo-structure and far again in ternary complex-structure), was reported for tomato and human GSNOR (Sanghani et al., 2002; Kubienova et al., 2013), but its function in the catalytic cycle is still an open issue. This alternate zinc ion positioning is not observed in the four subunits of apo-CrGSNOR1 structure (A, B, D and E) where Glu71 participates in metal coordination (Supplemental Figure 5D) and two subunits of the NAD^+^-CrGSNOR1 structure (C and F) where Glu71 is at a significantly higher distance than the other ligands (Supplemental Figure 5B). Interestingly, in four out of six subunits of the holo-form (A, C, E, and F) no water molecule was observed in close proximity to the catalytic zinc ion as found in other GSNOR structures. By contrast, in B and D subunits a water molecule is located at about 3 Å from the zinc ion at the opposite side with respect to Glu71 (Figure 4F). In the apo-structure, this water molecule always participates in metal coordination being hydrogen-bonded to Thr50 and Tyr96 (distance ranging from 4.2 to 5.8 Å; Supplemental Figure 6A). In all subunits of NAD^+^-CrGSNOR1 structure, the hydroxyl group of Tyr96 is rotated compared to the apo-form (Supplemental Figure 6B) and is not able to interact with the water molecule that partially loses its stabilization. Differently, Arabidopsis, tomato and human holo-structures always show a water molecule in the proximity of the catalytic zinc ion and the rotation of the conserved Tyr96 is not observed. When present, the water molecule bridges the zinc ion and the nicotinamide ring of the cofactor, which lies at about 5.0 Å from the catalytic metal (Figure 4F).

**Figure 4.**
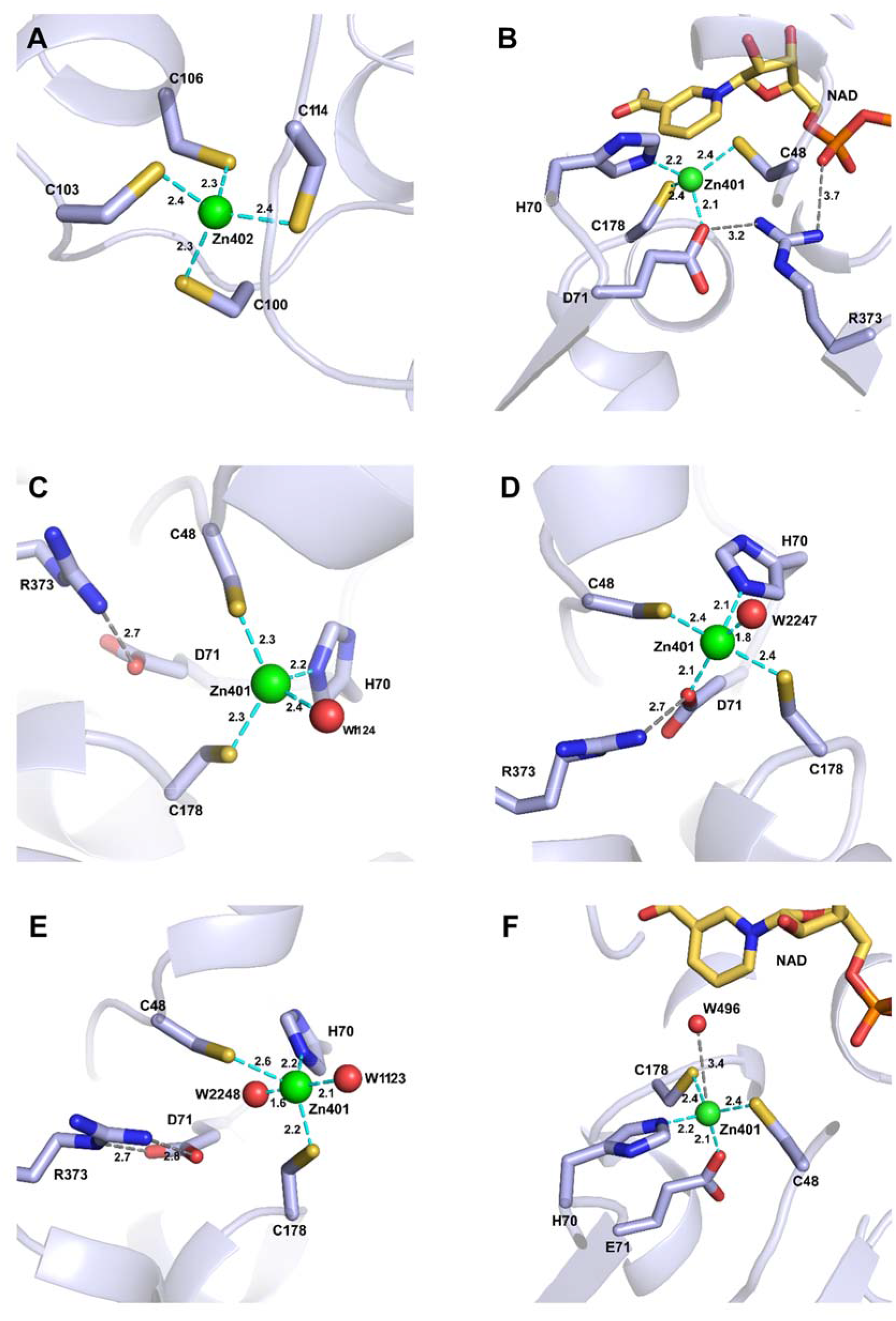
Coordination environment of the zinc ions in apo- and NAD^+^-CrGSNOR1. (**A**) The structural zinc ion is coordinated by four cysteine residues (100,103, 106 and 114) in all subunits of both enzyme forms. (**B**) The catalytic zinc ion is coordinated with a tetrahedral geometry by two cysteines (48 and 178), His70, and Glu71 in all subunits of NAD^+^-CrGSNOR1. Glu71 also forms a salt-bridge with Arg373, which in turn interacts with the phosphate groups of the cofactor. (**C**) In F subunit of apo-CrGSNOR1, the catalytic zinc ion is coordinated with a distorted tetrahedral geometry by Cys48, Cys178, His70, and a water molecule. Glu71 is uniquely involved in a salt-bridge with Arg373. (D) In A, B, D and E subunits of apo-CrGSNOR1, the catalytic zinc ion is coordinated by five ligands comprising Cys48, Cys178, His70, Glu71 and a water molecule. Glu71 keeps its interaction with Arg373. (**E**) In the C subunit of apo-CrGSNOR1, the catalytic zinc ion is coordinated by five ligands comprising Cys48, Cys178, His70, and two water molecules. Glu71 is uniquely involved in double salt-bridge with Arg373. (**F**) In B and D subunits of NAD^+^-CrGSNOR1, in close proximity to the catalytic zinc ion coordinated with a tetrahedral geometry by Cys48, Cys178, His70, and Glu71, a water molecule is observed.

**Figure 5.**
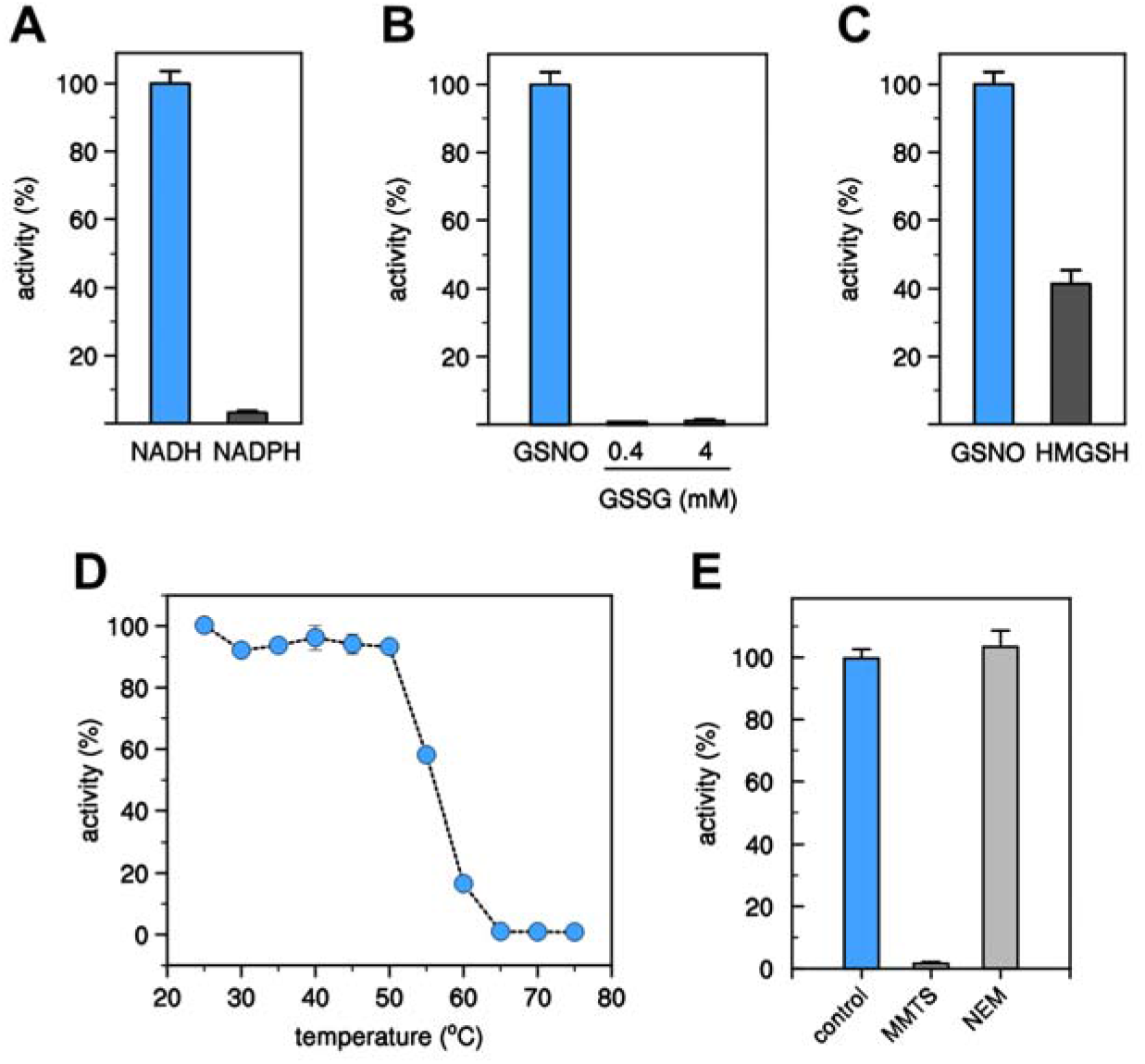
Kinetic analysis of CrGSNOR1. (**A**) Cofactor specificity of CrGSNOR1 activity in the presence of GSNO. The activity of CrGSNOR was evaluated in the presence of 0.4 mM GSNO using 0.2 mM NADH (white bar) or 0.2 mM NADPH (black bar) as cofactor. Data are represented as mean ± SD (n = 3). (**B**) Substrate specificity of CrGSNOR1 activity in the presence of NADH. The activity of CrGSNOR was evaluated in the presence of 0.2 mM NADH using 0.4 mM GSNO (white bar) or GSSG (0.4 or 4 mM, black bar) as substrate. Data are represented as mean ± SD (n = 3). (**C**) Activity of CrGSNOR1 as reductase or dehydrogenase. The activity of CrGSNOR was evaluated using GSNO (white bar) or HMGSH (black bar) as described in the Experimental section. Data are represented as mean ± SD (n = 3). For panels **A**-**C**, the NADH-dependent GSNO reduction of CrGSNOR1 (25 nM) was set to 100% (36.9 ± 2.9 μmol/min/mg). (**D**) Thermal stability of CrGSNOR1. The protein was incubated for 30 min at variable temperatures and after incubation, the remaining activity was measured. Data are represented as mean ± SD (n = 3). (**E**) Inactivation treatments of CrGSNOR1 with MMTS or NEM. CrGSNOR1 was incubated for 30 min in the presence of 1 mM MMTS or 1 mM NEM (gray bars). Data are represented as mean ± SD (n = 3) of control activity measured after protein incubation in the presence of buffer alone (light-blue bar).

The active site is located between the catalytic and cofactor-binding domains (Figure 3B) and is formed by several loops including Gly56-Glu61, Pro109-Val117, Ile93-Gln97, Phe144-Thr147, Ala286-Trp290 and Ile295-Gln303, and the α-helix Arg118-Lys123 (Figure 2). These portions contain residues involved in the binding of the substrate HMGSH in HsGSNOR (Engeland et al., 1993; Estonius et al., 1994; Sanghani et al., 2002). Most of these residues are conserved among different GSNORs (Figure 2), except for Gln112, Tyr140, and Lys284 (HsGSNOR numbering), which are replaced in CrGSNOR1 by Val115, Phe144, and Arg288, respectively (Figure 2). Within the substrate-binding site of NAD^+^-CrGSNOR1 structure (chains A-F), we observed a PEG molecule from the crystallization medium that had a different length in the diverse chains. The terminal hydroxyl group of PEG is located at more than 5.5 Å from the catalytic zinc ion and does not contribute to its coordination as observed for the hydroxyl group of HMGSH in HsGSNOR ternary complex (Sanghani et al., 2002). Hydrogen bonds with Tyr96, Gln97, NAD^+^, and several water molecules stabilize PEG (Supplemental Figure 4B). The rotation of Tyr96 side chain with respect to the position in the apo-structure is required for PEG accommodation into the substrate-binding site. An equivalent rotation is not observed in the HMGSH binding to HsGSNOR.

### Biochemical features of recombinant CrGSNOR1

Purified recombinant CrGSNOR1 was assayed for its ability to catabolize GSNO. The enzyme-catalyzed GSNO degradation in the presence of NADH displaying a linear relationship with protein concentrations (Supplemental Figure 7). By contrast, its activity was almost undetectable when NADPH replaced NADH (Figure 5A). Likewise, no activity was observed by replacing GSNO with GSSG (Figure 5B). These results indicate that CrGSNOR1 activity strictly depends on NADH and GSNO. GSNOR from diverse sources was originally found to catalyze the oxidation of S-(hydroxymethyl)glutathione (HMGSH) in the presence of NAD^+^ (Holmquist and Vallee, 1991; Liu et al., 2001; Sanghani et al., 2002; Kubienova et al., 2013; Matamoros et al., 2020). The Chlamydomonas GSNOR1 enzyme was also able to catalyze the NAD-dependent oxidation of HMGSH but with a 2.5-fold lower efficiency compared to the GSNO degrading activity (Figure 5C).

Kinetic analyses were performed on the NADH-dependent GSNO reducing activity of CrGSNOR1 using either GSNO or NADH as variable substrates and the kinetic parameters were determined by non-linear regression analysis (Supplemental Figure 8A and 8B). When the initial rates were plotted as a function of substrate concentration, responses were hyperbolic allowing apparent kinetic parameters to be calculated. The apparent Michaelis-Menten constants (*K*’_m_) measured at saturating concentrations of the non-varied substrate were 24.9 ± 1.5 μM for GSNO and 14.3 ± 2.1 μM for NADH and the apparent turnover numbers (*k*’_cat_) were 26.6 ± 2.5 sec^−1^ (GSNO) and 27.0 ± 0.6 sec^−1^ (NADH). The calculated catalytic efficiencies (*k*’_c_a_t_/*K*’_m_) of the reaction were ~1.07 x 10^6^ M^−1^ s^−1^ (GSNO) and ~1.86 x 10^6^ M^−1^ s^−1^ (NADH). These values are comparable to previously characterized plant GSNORs although kinetic properties slightly differ for CrGSNOR1 with a ~2–3-fold higher substrate/cofactor affinities and ~4–5-fold lower turnover numbers (Kubienova et al., 2013; Guerra et al., 2016; Ticha et al., 2017; Matamoros et al., 2020).

After establishing the biochemical properties of recombinant CrGSNOR1, we analyzed its sensitivity to thermal denaturation as carried out with Chlamydomonas protein extracts. The thermal stability of recombinant CrGSNOR1 was investigated by following the residual GSNOR activity after 30 min incubation at different temperatures (Figure 5D). The enzyme showed a relatively high degree of thermostability, retaining maximal activity in the 25-50 °C range. Exposure to higher temperatures led to a rapid protein inactivation being complete at temperatures above 65 °C. These observations correlate with the thermal sensitivity of the NADH-dependent activity measured in algal protein extracts (Figure 1d). T_50_, the temperature at which 50% of the activity is retained after 30 min incubation, was found to be ~56 °C, a value strikingly similar to other plant GSNORs (Kubienova et al., 2013; Ticha et al., 2017). The effect of temperature on CrGSNOR1 stability was also evaluated by following the turbidity at 405 nm, which represents an optical measurement for protein denaturation/aggregation (Supplemental Figure 8C). Consistent with activity measurements, CrGSNOR1 remained fully stable when incubated at 25 °C, whereas it started to aggregate immediately after incubation at 75 °C, reaching maximal turbidity after 10 min. At 55 °C, the aggregation kinetic proceeded more slowly and half-maximal turbidity was reached after 30 min.

To further extend the comparison between the recombinant protein and the NADH-dependent enzymatic system from algal protein extracts, we examined the sensitivity of CrGSNOR1 to MMTS and NEM. Exposure of CrGSNOR1 to MMTS resulted in a complete inactivation of the enzyme, while NEM did not affect catalysis (Figure 5E). This observation is in agreement with the catalytic effect of these two thiol-modifying compounds on algal protein extracts where NADH-dependent activity was only affected in the presence of MMTS (Figure 1F). In protein extracts, however, the MMTS-dependent inactivation was only partial and this might be due to its reaction with thiol-containing proteins other than GSNORs. These results also indicate that CrGSNOR1 activity has a dissimilar response to MMTS and NEM, likely residing on the reactivity of MMTS with both solvent accessible and zinc-coordinating cysteines (D’Ordine et al., 2012) thus affecting protein catalysis and/or structural stability.

### Cysteine conservation and thiol reactivity in CrGSNOR1

CrGSNOR1 is a cysteine-rich enzyme as it contains sixteen cysteines that correspond to 4.2% of the total amino acid content (Supplemental Figure 1). Among GSNORs from different species, nine out of sixteen cysteines are fully conserved comprising Cys48/Cys178 (coordination of the catalytic zinc atom), Cys100/Cys103/Cys106/Cys114 (coordination of the structural zinc atom) and Cys11 (except in bacterial GSNORs), Cys174 (except in *C. elegans*) and Cys272 (Figure 2 and Supplemental Figure 2). The remaining seven Cys are randomly conserved with Cys95/Cys285/Cys371/Cys374 only present in the green lineage with the exceptions of Cys95 absent in pea GSNOR, Cys374 absent in the tomato and *Lotus japonicus* GSNOR1, and Cys285/Cys371 present in Synechocystis/yeast GSNOR, respectively. Cys244 is conserved in algae and most animals and bacteria (Figure 2 and Supplemental Figure 2). Despite the high Cys content, we found that CrGSNOR1 only contains two solvent accessible/reactive cysteine thiols as assessed by DTNB-based thiol titration (2.0 ± 0.3 free thiols per subunit).

In order to confirm the number of accessible/reactive free thiols and establish their position, we analyzed the protein by matrix-assisted laser desorption ionization time-of-flight mass spectrometry (MALDI-TOF MS) following alkylation treatment in the presence of maleimide derivatives. Preliminary NEM-based alkylation experiments suggested that CrGSNOR1 was mainly di-alkylated (Supplemental Figure 9). Nevertheless, the low mass shift induced by NEM (+125 Da per alkylated cysteine) precluded a clear separation of the different alkylated forms of CrGSNOR1 at the protein level. Therefore, NEM was replaced by Biotin-maleimide as it exhibits the same maleimide reactive group but allows better separation of the different protein species by generating a +451 Da mass shift per alkylated cysteine. As shown in Figure 6, CrGSNOR1 underwent a near complete di-alkylation after 30 min incubation and longer incubation showed no further significant peaks. These data are consistent with the two accessible/reactive cysteine thiols determined by DTNB assay. Subsequently, we identified the alkylated cysteines by peptide mass fingerprinting of CrGSNOR1 treated with Biotin-maleimide for 20 min. This incubation time was selected to generate partial mono- and di-alkylated species of CrGSNOR1. By comparing MALDI-TOF spectra obtained after trypsin digestion of untreated or Biotin-maleimide-treated CrGSNOR1, we identified Cys244 and Cys371 as alkylated residues (Figure 7). Taking advantage of the presence of a biotin moiety, we also performed an enrichment of alkylated peptides using monomeric avidin as previously described in (Pérez-Pérez et al., 2017) and we confirmed the alkylation of Cys371 (Supplemental Figure 10) while peptides containing Cys244 were not recovered likely due to its weak propensity to ionize under MALDI ionization conditions. Altogether, mass spectrometry analyses are consistent with the structural features of CrGSNOR1. Alkylation of Cys371 agrees with the high accessible surface area (ASA) calculated from the structure, ranging from 14 Å^2^ to 31 Å^2^ in different chains of the asymmetric unit. Similarly, Cys244 has an accessibility in the 29-31 Å^2^ range, supporting its reactivity towards maleimide. The structure of the apo-form revealed that Cys272 is also exposed to the solvent (ASA 16-22 Å^2^) but no alkylation was observed (Supplemental Figure 11). This lack of reactivity may depend on the orientation of its thiol group toward a hydrophobic cavity (formed by Val187, 193, 201, 207, 211 and 296, Ala186, Gly208 and Phe270) that likely hampers reaction with maleimide derivatives. Conversely, the cofactor binding makes Cys272 completely buried in the holo-form.

**Figure 6.**
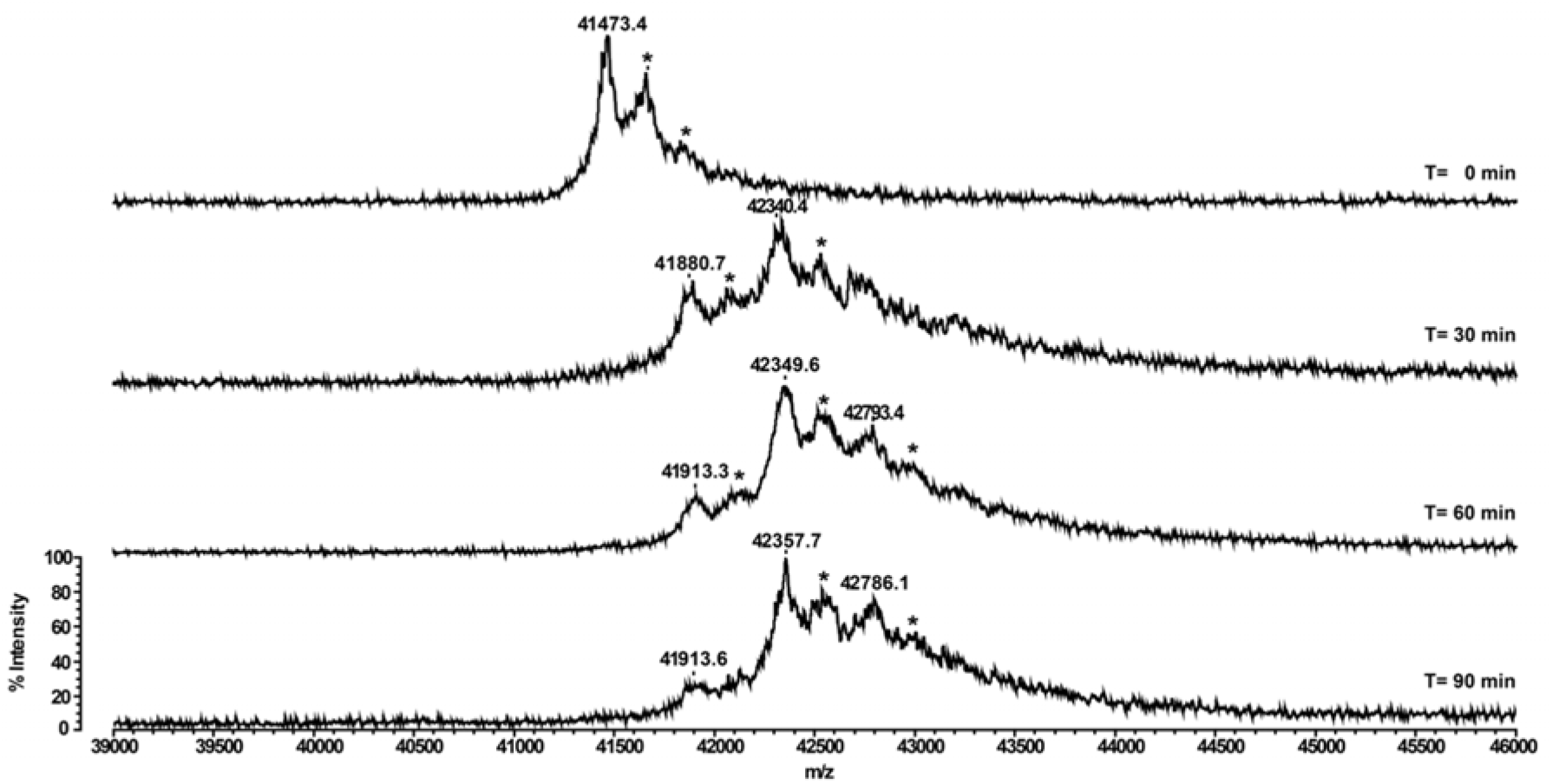
Time-dependent mass spectrometry analyses of CrGSNOR1 treated with Biotin-maleimide. Recombinant CrGSNOR1 was incubated in the presence of 1 mM Biotin-maleimide. At indicated time points, protein samples were withdrawn and analyzed by MALDI-TOF MS to assess the number of alkylated cysteines. For each alkylated cysteine, the molecular mass of CrGSNOR1 is shifted by +451 Da compared to the native protein (41473.4 Da). Peaks highlighted by an asterisk correspond to the protein-matrix (sinapinic acid) adduct. The y-axis is equal for all mass spectra acquired at times 0, 30, 60, and 90 min, and only indicated in the bottom spectrum.

**Figure 7.**
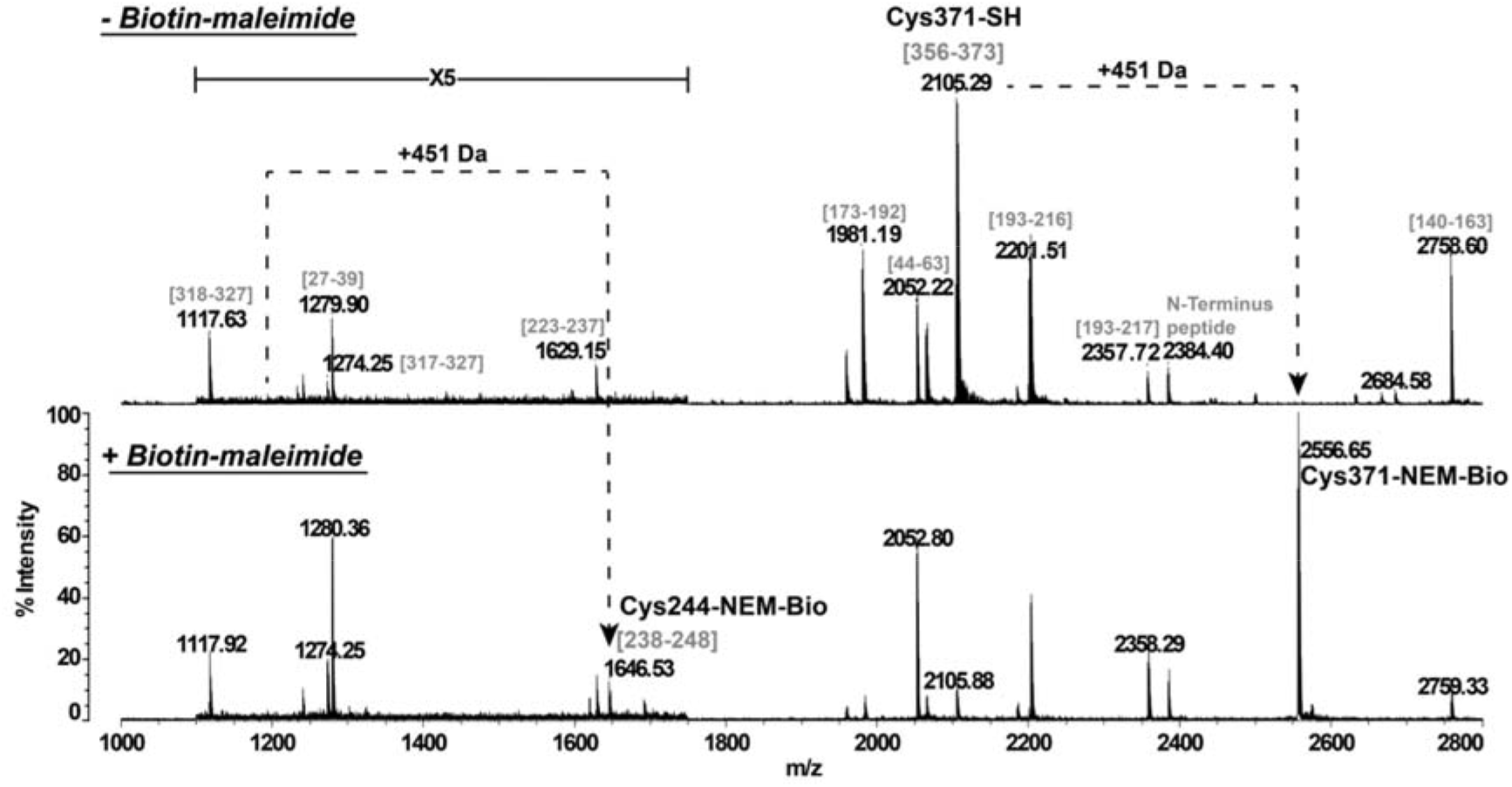
Peptide mass fingerprinting of untreated or Biotin-maleimide-treated CrGSNOR1. Recombinant CrGSNOR1 was incubated in the presence of 1 mM Biotin-maleimide for 20 min and then trypsin digested. The peptide mixture was analyzed by MALDI-TOF MS. Sequence of peptides belonging to CrGSNOR1 is indicated in brackets (numbering according to Figure 2). Cysteines modified by Biotin-maleimide are annotated with the mention “NEM-Bio” and the peak corresponding to the peptide sequence [1-12] of CrGSNOR1 is indicated as N-terminus peptide as it is fused with the 7xHis affinity purification tag.

### CrGSNOR1 has limited sensitivity to S-nitrosylation and remains unaffected by oxidative treatments

Thiol-modifying treatments suggested that CrGSNOR1 contains cysteine(s) that might be prone to oxidative modifications that may affect enzyme catalysis. To investigate the sensitivity of CrGSNOR1 to physiological thiol-based modifications, we measured protein activity upon treatments with different molecules that specifically induce cysteine oxidation. As shown in Figure 8A, diamide (TMAD) and hydrogen peroxide (H_2_O_2_) did not significantly alter CrGSNOR1 activity even at a high concentration (1 mM). Moreover, circular dichroism (CD) spectra of apo-CrGSNOR1 before and after treatment with H_2_O_2_ are substantially superimposable, ruling out a significant variation of secondary structure upon the oxidative treatment (Supplemental Figure 12 and Supplemental Table 2). Previous studies reported that plant GSNORs undergo S-nitrosylation with consequent inhibition of nearly all isoforms with the exception of GSNOR1 and GSNOR2 from *Lotus japonicus* (Frungillo et al., 2014; Cheng et al., 2015; Guerra et al., 2016; Ticha et al., 2017; Zhan et al., 2018; Chen et al., 2020; Matamoros et al., 2020; Zhang et al., 2020). In order to examine whether S-nitrosylation can regulate CrGSNOR1, the purified enzyme was exposed to different types of NO-donors. Nitrosylation reactions were induced chemically with the NO-releasing compound SNAP or with GSNO that acts as a trans-nitrosylating agent (Askew et al., 1995). In the presence of GSNO, the activity of CrGSNOR1 remained unaffected (Figure 8B) while we observed a partial and reversible inhibition in the presence of SNAP (Figure 8B and 8C). To assess the S-nitrosylation status of CrGSNOR1, we applied the biotin switch technique (BST) on the GSNO- and SNAP-treated enzyme. Surprisingly, we observed a positive nitrosylation signal following exposure to both nitrosylating agents (Figure 8D and 8E), indicating that either GSNO or SNAP can induce cysteine S-nitrosylation but only the latter was found to affect, though partially, CrGSNOR1 catalysis.

**Figure 8.**
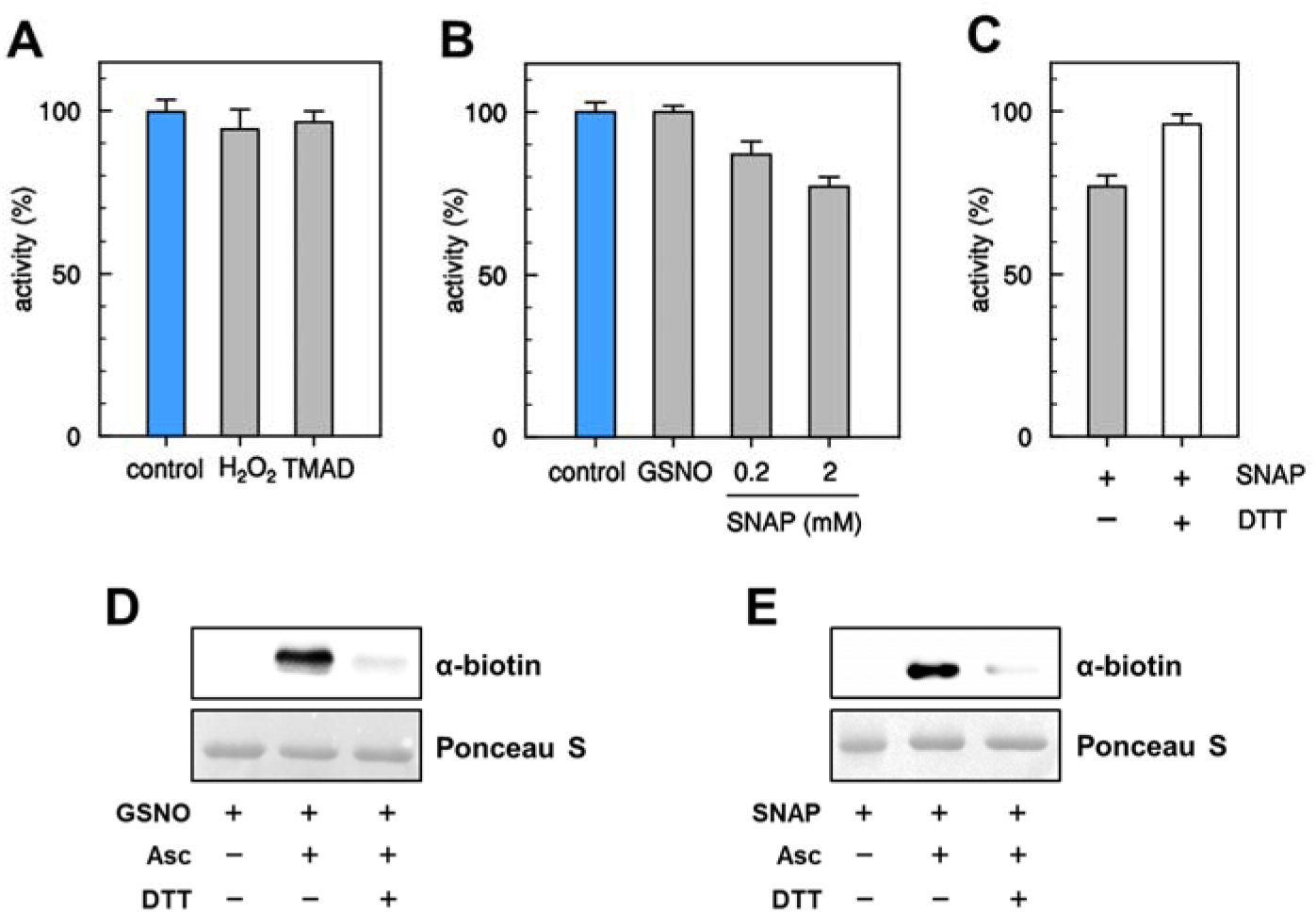
Effects of oxidizing and nitrosylating agents on CrGSNOR1. **(A)** Incubation of CrGSNOR1 with oxidizing agents. CrGSNOR1 was incubated for 30 min in the presence of 1 mM H_2_O_2_ or diamide (TMAD) (grey bars). Data are represented as mean ± SD (n = 3) of control activity measured after 30 min incubation in the absence of oxidizing agents (light blue bar). **(B)** Incubation of CrGSNOR1 with the nitrosylating agents GSNO and SNAP. CrGSNOR1 was incubated for 30 min in the presence of GSNO (2 mM) or SNAP (0.2 or 2 mM). Data are represented as mean ± SD (n = 3) of control activity measured after 30 min incubation in the absence of DEA-NO (white bar). **(C)** The reversibility of CrGSNOR1 inactivation by SNAP (2 mM, black bar) was assessed by incubation in the presence of 20 mM DTT (white bars). Data are represented as mean ± SD (n = 3) of control activity (see panel B). **(D-E)** S-nitrosylation of CrGSNOR1. CrGSNOR1 was treated for 30 min in the presence of 2 mM GSNO (D) or 2 mM SNAP (E) and nitrosylation was visualized using the biotin-switch technique followed by anti-biotin western blots as described in Material and Methods. For both panels, the red-ponceau (ponceau) staining of the membranes shows almost equal loading in each lane. Asc, ascorbate.

## DISCUSSION

Over the last decades, NO signaling has emerged as a fundamental process by which photosynthetic organisms including unicellular algae, regulate different aspects of cell metabolism (Zaffagnini et al., 2016; Del Castello et al., 2019; Kolbert et al., 2019). Characterization of the mechanisms regulating NO homeostasis and NO-dependent signaling pathway, is of striking importance in microalgae considering their biotechnological potential for the bio-production of drugs, energy and food (Wijffels et al., 2013; Scaife et al., 2015). Recent efforts have set the foundations for the use of *Chlamydomonas* and other unicellular phototrophs as molecular chassis exploitable in the next synthetic biology-driven green revolution (Scaife and Smith, 2016; Crozet et al., 2018; Vavitsas et al., 2019). NO metabolism is of particular importance in these organisms living in liquid micro-oxic environments, where the fermentative metabolism and the Hemoglobin-NO cycle are important players in cellular bioenergy (Hemschemeier et al., 2013; Becana et al., 2020). In *Chlamydomonas*, the biological function of NO relates to responses to nitrogen and sulfur starvation, hypoxia/anoxia, high light and light to dark transitions (Hemschemeier et al., 2013; Wei et al., 2014; Berger et al., 2016; Pokora et al., 2017; De Mia et al., 2019; Kuo et al., 2020). In general, protein S-nitrosylation acts as the major mechanism propagating NO-dependent biological signaling and it can modulate protein function by altering enzymatic activity and/or protein structure (Zaffagnini et al., 2016; Feng et al., 2019; Stomberski et al., 2019; Zaffagnini et al., 2019). Noteworthy, while NO-dependent biological pathways have been uncovered, very little is known about how microalgae control nitrosothiol homeostasis through NO catabolism.

Considering the primary role of GSNO as a trans-nitrosylating agent, the redox systems involved in GSNO catabolism are fundamental to control the intracellular levels of this low-molecular weight nitrosothiol and consequently, the extent of protein S-nitrosylation. In this work, we observed that *Chlamydomonas* protein extracts contain two distinct systems catalyzing GSNO reduction using NADPH or NADH and having different sensitivities to thiol-modifying compounds and thermal denaturation. Based on cofactor specificity and biochemical properties, we can hypothesize that the NADPH-dependent activity might primarily involve thiol-disulfide exchanges mediated by TRX or GRX (Sengupta and Holmgren, 2012; Ren et al., 2019). Indeed, these enzymes are thermostable and their activity is inhibited by irreversible alkylation (Lemaire et al., 2000; Marchand et al., 2019). Similar properties are also found in glutathione reductases but these enzymes cannot use GSNO as a substrate ((Becker et al., 1995) and authors’ personal communication). Other NADPH-dependent GSNO-reducing activities might be involved such as carbonyl reductase 1 and aldo-keto reductase family 1 member A1 identified in human (Bateman et al., 2008; Stomberski et al., 2019) and for which orthologs are present in Chlamydomonas genome (data not shown). The NADH dependent activity observed in Chlamydomonas protein extracts is most likely dependent on GSNOR since its sensitivity to thiol modifying agents and thermal denaturation is comparable to purified Chlamydomonas GSNOR1, which was found to strictly depend on GSNO and NADH (Figure 1 and Figure 5). This is further supported by an overall conservation of the three-dimensional structure of apo- and holo-CrGSNOR1 compared to other structurally known GSNORs. Despite this global structural homology, we observed that the coordination sphere of the catalytic zinc ion shows a high variability (Figure 4B-F), while the tetrahedral thiolate-geometry (S4) coordination of the structural zinc ion is perfectly conserved (Figure 4A). When NAD^+^ is bound to CrGSNOR1, the zinc atom is mainly coordinated by four conserved residues (Cys48, His70, Glu71 and Cys178; Figure 2 and Figure 4B) as in human, tomato, and Arabidopsis GSNORs. However, in two out of six subunits, the metal stabilization by Glu71 appears weaker with a distance above 3.5 Å. Indeed carboxylate groups can show a wide range of metal-ligand distances up to 4.5 Å (Maret and Li, 2009). Differently, in the apo-structure the catalytic zinc is stabilized by four or five ligands involving Cys48, His70, Cys178, and Glu71 replaced or accompanied by one or two water molecules (or hydroxide ions; Figure 4C-F). This expansion to a penta-coordination sphere, already reported for other zinc-containing proteins (*e.g*. adenosine deaminase (Wilson and Quiocho, 1993); astacin (Gomis-Ruth et al., 1993)) or temporarily occurring in catalytic zinc-sites to accommodate the substrate (Holmes and Matthews, 1981; McCall et al., 2000; Daniel and Farrell, 2014), was not observed in other structurally known GSNORs. The functional role of this increased dynamicity of the catalytic zinc in the algal enzyme possibly due to steric and stabilizing electrostatic interactions from the secondary coordination sphere, remains to be established.

GSNOR is typically defined as a cysteine-rich protein containing 14 to 16 Cys residues (Figure 2 and Supplemental Figure 2). Consistently, CrGSNOR1 contains 16 Cys of which only Cys244 and Cys371 were found to be solvent-exposed and reactive towards maleimide derivatives, although alkylation did not affect activity, in sharp contrast with AtGSNOR whose activity is strongly inhibited after exposure to alkylating agents (Kovacs et al., 2016). While CrGSNOR1 is resistant to NEM, MMTS-dependent conjugation causes a rapid inactivation of the enzyme, which is likely ascribed to the ability of MMTS to alter zinc(s)-coordination with consequent protein inactivation. The response of CrGSNOR1 to thiol-based modifications reflects dissimilarities with other plant GSNORs (Lindermayr, 2017; Jahnova et al., 2019). Recent studies showed that GSNOR from several land plants was inhibited by both H_2_O_2_-dependent oxidation and S-nitrosylation (Frungillo et al., 2014; Cheng et al., 2015; Ticha et al., 2017; Zhan et al., 2018; Zhang et al., 2020). Lindermayr and colleagues identified the catalytic zinc-coordinating cysteine residues (Cys47 and Cys177) as primary targets of H_2_O_2_-mediated oxidation in AtGSNOR with consequent zinc ion release and disruption of the catalytic site (Kovacs et al., 2016). Although these cysteines are fully conserved in CrGSNOR1, oxidizing compounds such as diamide and H_2_O_2_ did not alter protein activity and folding (Figure 8 and Supplemental Figure 12). This suggests that CrGSNOR1 does not contain oxidation-prone zinc-binding cysteine(s), likely due to protection by a highly stable coordination in the algal enzyme. CrGSNOR1 was found to undergo S-nitrosylation but without any significant effect on enzyme activity (Figure 8), as previously observed for GSNORs from *Lotus Japonicus* (Matamoros et al., 2020). By contrast, AtGSNOR was shown to undergo S-nitrosylation on Cys10, Cys271, and Cys370, leading to inhibition of the enzyme through a 2-fold decrease of both the affinity towards GSNO and the turnover number (Guerra et al., 2016). These three residues are fully conserved in CrGSNOR1 (Figure 2 and Figure 9A), but alterations in cysteine microenvironments and local folding might explain the limited responsiveness of CrGSNOR1 to S-nitrosylation or other thiol modifications.

**Figure 9.**
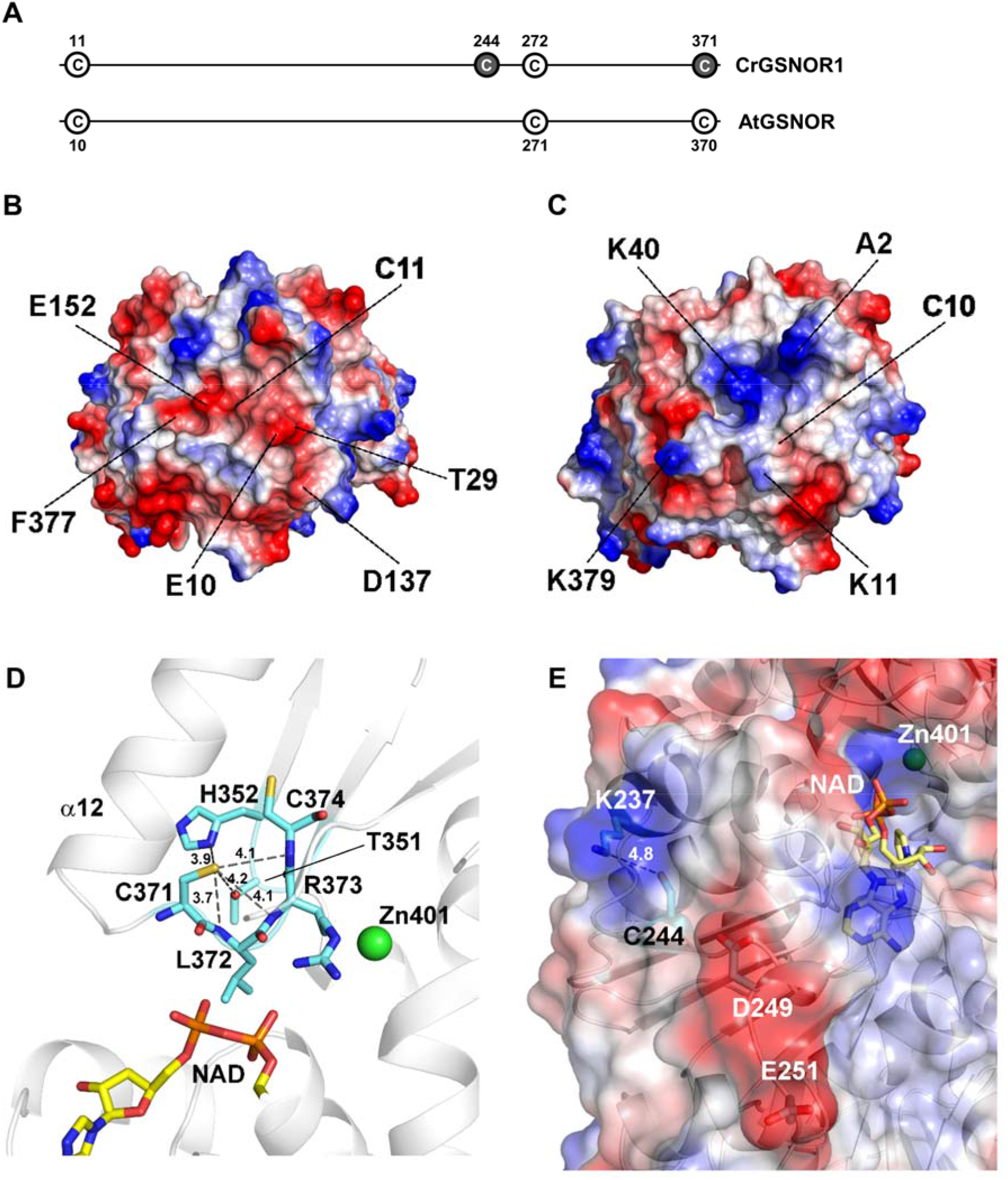
Microenvironment of thiol-modified cysteine residues in algae and/or plant GSNORs. **(A)** Conservation of modified cysteine residues in CrGSNOR1 (alkylated Cys dark circle) and AtGSNOR (S-nitrosylated Cys white circle). Cys11 and Cys272 not modified in CrGSNOR1 are reported as a white circle. Electrostatic surface potential in the region surrounding Cys11 in CrGSNOR1 **(B)** and Cys10 in AtGSNOR **(C)**. Specific residues determining the larger differences in surface potential between the two enzymes are indicated. The electrostatic surface potential ranges between −60 (red) and 60 (blue). **(D)** CrGSNOR1 Cys371 lies in a loop that follows helix α12. Its thiol group is hydrogen-bonded to the side chains and backbone nitrogen atoms of several surrounding residues. Cys371 overlooks the catalytic cavity containing the zinc ion and the cofactor. **(E)** The thiol group of CrGSNOR1 Cys244 is hydrogen-bonded to the amino side chain group of Lys237, which determines a positive surface electrostatic potential. The carboxylic group of Asp249 and Glu251 determine a negative region on the other side of Cys244 thiol group.

In CrGSNOR1, Cys11 is barely accessible, as in AtGSNOR, and not reactive toward alkylating reagents. Structural comparison between the two enzymes unveiled that despite a generally high backbone similarity, Cys11 in CrGSNOR1 is surrounded by specific residues (*i.e*. Glu10, Thr29, Asp137, Glu152 and the carboxyl group of C-terminal Phe377) that determine a negative electrostatic surface potential (Figure 9B). These residues are not conserved in AtGSNOR1 (Figure 2) where Cys10 is rather surrounded by positive charges (Figure 9C) due to the N-terminal groups of Ala2, Lys11, Lys40 and Lys379 (the two latter not conserved in CrGSNOR1, Figure 2). The differences between the microenvironment of Cys11/Cys10 in the two enzymes could be the basis for the lack of reactivity of this residue in CrGSNOR1. *In vivo*, nitrosylation of AtGSNOR Cys10 occurs in response to hypoxia and promotes degradation of the enzyme by selective autophagy (Zhan et al., 2018). Recently, the non-canonical catalase CAT3 was shown to transnitrosylate AtGSNOR1 at Cys 10, a process strictly dependent on CAT3 Cys343 (Chen et al., 2020). This mechanism is unlikely to occur in Chlamydomonas since the previously cited cysteine is not conserved in the unique algal redox regulated catalase (Shao et al., 2008; Michelet et al., 2013).

As observed for Cys370 in AtGSNOR (Xu et al., 2013), CrGSNOR1 Cys371 is exposed to the solvent, although its accessibility decreases from the apo-to the holo-form (31-14 Å^2^ to 15-5 Å^2^), and undergoes maleimide-dependent alkylation. Nevertheless, Cys371 alkylation has no effect on CrGSNOR1 activity. In both enzymes, Cys371/Cys370 lies in a loop that follows helix α12 and its thiol group forms various hydrogen bonds with surrounding residues (Figure 2 and Figure 9D). This region is characterized by high mobility expressed by backbone thermal parameters (B factors) larger than the average value for the whole protein (Supplemental Figure 13 and Supplemental Table 1). However, in AtGSNOR, helix α12 is 3-4 residues shorter compared to CrGSNOR1 thereby decreasing the structural constraints due to a rigid secondary structure and increasing the probability that a redox modification of the cysteine could induce a local conformational rearrangement affecting the catalytic activity. Indeed, this residue overlooks the cofactor binding pocket and lies at about 11-12 Å from the catalytic zinc ion (Figure 9D). As observed for Cys371, also Cys272 (Cys271 in AtGSNOR) is solvent-accessible in CrGSNOR1. However, this residue becomes completely buried when NAD(H) is bound to the enzyme on the other side of the cofactor-binding pocket. By comparing the microenvironment of this residue, we observed that the region surrounding Cys272/Cys271 is structurally conserved between algal and plant enzymes. However, this residue is not modified by alkylation in CrGSNOR1 and we can thus suppose that physiological oxidative molecules cannot alter its redox state.

The other solvent-exposed cysteine targeted by maleimide alkylation in the Chlamydomonas enzyme, Cys244, is not conserved in GSNORs from land plants while it is present in other organisms including *V. carteri*, human, mouse, *C. elegans*, and prokaryotes such as *E. coli* and *Synechocystis sp*. PCC6803 (Figure 2 and Supplemental Figure 2). CrGSNOR1 Cys244 is hydrogen-bonded to Lys237 (Figure 9E) and the presence of a positive region determined by Lys237 side chain in close proximity to the Cys244 thiol group and of a larger negative region due to Asp249 and Glu251, determine a specific microenvironment that can facilitate a proper binding of oxidative molecules (*e.g*. NO or GSNO). Nevertheless, alkylation of this residue did not alter the catalytic functioning of the enzyme, suggesting that modification of this residue is unlikely to control CrGSNOR1.

Finally, we demonstrated that CrGSNOR1 has a null or limited sensitivity to thiol-based oxidative modifications compared to Arabidopsis GSNOR (Guerra et al., 2016; Kovacs et al., 2016) and other homologues from land plants (Ticha et al., 2017) with the notable exception of *Lotus Japonicus* GSNORs (Matamoros et al., 2020). Deep analyses of the crystallographic structure of CrGSNOR1 revealed structural features likely responsible for the difference in Cys reactivity compared to plant enzymes. Indeed, cysteine reactivity does not reflect the absolute solvent accessibility of the residue and is also influenced by the cysteine microenvironment or local folding. Specifically, these structural features can (i) protect the residues from oxidative attacks, (ii) hamper proper allocation of oxidative compounds, and (iii) limit conformational changes that might directly affect protein catalysis or favor the redox sensitivity of other cysteine residues. The limited sensitivity of CrGSNOR1 to redox post-translational modifications suggests that regulation of NO signaling in algae may operate through other mechanisms including regulation of GSNOR by other modifications or by regulation of protein abundance and gene expression. GSNOR may also be constitutively active in algae, regulation of nitrosothiols abundance being controlled by other NO degrading enzymes or at the level of NO production. These differences are likely linked to distinct requirements for regulation of NO metabolism between land plants and algae.

## ACKNOWLEDGMENTS

We gratefully acknowledge Elettra and the European Synchrotron Radiation Facility (ESRF) for allocation of beam time. SF and GF S.F. thanks the Consorzio Interuniversitario di Ricerca in Chimica dei Metalli nei Sistemi Biologici (CIRCMSB). This work was supported by University of Bologna Alma Idea 2017 Program (to MZ); CNRS Sorbonne Université, Agence Nationale de la Recherche Grant 17-CE05-0001 CalvinDesign (to CHM and SDL); LABEX DYNAMO (ANRLABX-011 to CHM, MDM, and SDL) and EQUIPEX CACSICE (ANR-11-EQPX-0008 to CHM and SDL), partly through funding of the Proteomic Platform IBPC (PPI). JR is supported by a PhD grant from the University of Bologna (PhD program in Cellular and Molecular Biology).

## Parsed Citations

Adams PD, Afonine PV, Bunkoczi G, Chen VB, Davis IW, Echols N, Headd JJ, Hung LW, Kapral GJ, Grosse-Kunstleve RW, McCoy AJ, Moriarty NW, Oeffner R, Read RJ, Richardson DC, Richardson JS, Terwilliger TC, Zwart PH (2010) PHENIX: a comprehensive Python-based system for macromolecular structure solution. Acta Crystallogr D Biol Crystallogr 66: 213–221

Ageeva-Kieferle A, Rudolf EE, Lindermayr C (2019) Redox-Dependent Chromatin Remodeling: A New Function of Nitric Oxide as Architect of Chromatin Structure in Plants. Front Plant Sci 10: 625

Airaki M, Sanchez-Moreno L, Leterrier M, Barroso JB, Palma JM, Corpas FJ (2011) Detection and quantification of S-nitrosoglutathione (GSNO) in pepper (Capsicum annuum L.) plant organs by LC-ES/MS. Plant Cell Physiol 52: 2006–2015

Askew SC, Butler AR, Flitney FW, Kemp GD, Megson IL (1995) Chemical mechanisms underlying the vasodilator and platelet antiaggregating properties of S-nitroso-N-acetyl-DL-penicillamine and S-nitrosoglutathione. Bioorg Med Chem 3: 1–9

Astier J, Kulik A, Koen E, Besson-Bard A, Bourque S, Jeandroz S, Lamotte O, Wendehenne D (2012) Protein S-nitrosylation: what’s going on in plants? Free Radic Biol Med 53: 1101–1110

Bai XG, Chen JH, Kong XX, Todd CD, Yang YP, Hu XY, Li DZ (2012) Carbon monoxide enhances the chilling tolerance of recalcitrant Baccaurea ramiflora seeds via nitric oxide-mediated glutathione homeostasis. Free Radic Biol Med 53: 710–720

Bateman RL, Rauh D, Tavshanjian B, Shokat KM (2008) Human carbonyl reductase 1 is an S-nitrosoglutathione reductase. J Biol Chem 283: 35756–35762

Becana M, Yruela I, Sarath G, Catalan P, Hargrove MS (2020) Plant hemoglobins: a journey from unicellular green algae to vascular plants. New Phytol

Becker K, Gui M, Schirmer RH (1995) Inhibition of human glutathione reductase by S-nitrosoglutathione. Eur J Biochem 234: 472–478

Begara-Morales JC, Sanchez-Calvo B, Chaki M, Valderrama R, Mata-Perez C, Padilla MN, Corpas FJ, Barroso JB (2016) Antioxidant Systems are Regulated by Nitric Oxide-Mediated Post-translational Modifications (NO-PTMs). Front Plant Sci 7: 152

Bellin D, Asai S, Delledonne M, Yoshioka H (2013) Nitric oxide as a mediator for defense responses. Mol Plant Microbe Interact 26: 271–277

Berger H, De Mia M, Morisse S, Marchand CH, Lemaire SD, Wobbe L, Kruse O (2016) A Light Switch Based on Protein S-Nitrosylation Fine-Tunes Photosynthetic Light Harvesting in Chlamydomonas. Plant Physiol 171: 821–832

Besson-Bard A, Pugin A, Wendehenne D (2008) New insights into nitric oxide signaling in plants. Annu Rev Plant Biol 59: 21–39

Bischof JC, He X (2005) Thermal stability of proteins. Ann N Y Acad Sci 1066: 12–33

Broniowska KA, Diers AR, Hogg N (2013) S-Nitrosoglutathione. Biochim Biophys Acta

Calatrava V, Chamizo-Ampudia A, Sanz-Luque E, Ocana-Calahorro F, Llamas A, Fernandez E, Galvan A (2017) How Chlamydomonas handles nitrate and the nitric oxide cycle. J Exp Bot 68: 2593–2602

Chen L, Wu R, Feng J, Feng T, Wang C, Hu J, Zhan N, Li Y, Ma X, Ren B, Zhang J, Song CP, Li J, Zhou JM, Zuo J (2020) Transnitrosylation Mediated by the Non-canonical Catalase ROG1 Regulates Nitric Oxide Signaling in Plants. Dev Cell 53: 444–457 e445

Chen VB, Arendall WB, 3rd, Headd JJ, Keedy DA, Immormino RM, Kapral GJ, Murray LW, Richardson JS, Richardson DC (2010) MolProbity: all-atom structure validation for macromolecular crystallography. Acta Crystallogr D Biol Crystallogr 66: 12–21

Chen X, Tian D, Kong X, Chen Q, E FA, Hu X, Jia A (2016) The role of nitric oxide signalling in response to salt stress in Chlamydomonas reinhardtii. Planta 244: 651–669

Cheng T, Chen J, Ef AA, Wang P, Wang G, Hu X, Shi J (2015) Quantitative proteomics analysis reveals that S-nitrosoglutathione reductase (GSNOR) and nitric oxide signaling enhance poplar defense against chilling stress. Planta 242: 1361–1390

Corpas FJ, Alche JD, Barroso JB (2013) Current overview of S-nitrosoglutathione (GSNO) in higher plants. Front Plant Sci 4: 126

Crozet P, Navarro FJ, Willmund F, Mehrshahi P, Bakowski K, Lauersen KJ, Perez-Perez ME, Auroy P, Gorchs Rovira A, Sauret-Gueto S, Niemeyer J, Spaniol B, Theis J, Trosch R, Westrich LD, Vavitsas K, Baier T, Hubner W, de Carpentier F, Cassarini M, Danon A, Henri J, Marchand CH, de Mia M, Sarkissian K, Baulcombe DC, Peltier G, Crespo JL, Kruse O, Jensen PE, Schroda M, Smith AG, Lemaire SD (2018) Birth of a Photosynthetic Chassis: A MoClo Toolkit Enabling Synthetic Biology in the Microalga Chlamydomonas reinhardtii. ACS Synth Biol 7: 2074–2086

D’Ordine RL, Linger RS, Thai CJ, Davisson VJ (2012) Catalytic zinc site and mechanism of the metalloenzyme PR-AMP cyclohydrolase. Biochemistry 51: 5791–5803

Daniel AG, Farrell NP (2014) The dynamics of zinc sites in proteins: electronic basis for coordination sphere expansion at structural sites. Metallomics 6: 2230–2241

De Mia M, Lemaire SD, Choquet Y, Wollman FA (2019) Nitric Oxide Remodels the Photosynthetic Apparatus upon S-Starvation in Chlamydomonas reinhardtii. Plant Physiol 179: 718–731

Del Castello F, Nejamkin A, Cassia R, Correa-Aragunde N, Fernandez B, Foresi N, Lombardo C, Ramirez L, Lamattina L (2019) The era of nitric oxide in plant biology: Twenty years tying up loose ends. Nitric Oxide 85: 17–27

Emsley P, Cowtan K (2004) Coot: model-building tools for molecular graphics. Acta Crystallogr D Biol Crystallogr 60: 2126–2132

Engeland K, Hoog JO, Holmquist B, Estonius M, Jornvall H, Vallee BL (1993) Mutation of Arg-115 of human class III alcohol dehydrogenase: a binding site required for formaldehyde dehydrogenase activity and fatty acid activation. Proc Natl Acad Sci U S A 90: 2491–2494

Estonius M, Hoog JO, Danielsson O, Jornvall H (1994) Residues specific for class III alcohol dehydrogenase. Site-directed mutagenesis of the human enzyme. Biochemistry 33: 15080–15085

Evans PR, Murshudov GN (2013) How good are my data and what is the resolution? Acta Crystallogr D Biol Crystallogr 69: 1204–1214

Feechan A, Kwon E, Yun BW, Wang Y, Pallas JA, Loake GJ (2005) A central role for S-nitrosothiols in plant disease resistance. Proc Natl Acad Sci U S A 102: 8054–8059

Feng J, Chen L, Zuo J (2019) Protein S-Nitrosylation in plants: Current progresses and challenges. J Integr Plant Biol 61: 1206–1223

Foster MW, Liu L, Zeng M, Hess DT, Stamler JS (2009) A genetic analysis of nitrosative stress. Biochemistry 48: 792–799

Frungillo L, Skelly MJ, Loake GJ, Spoel SH, Salgado I (2014) S-nitrosothiols regulate nitric oxide production and storage in plants through the nitrogen assimilation pathway. Nat Commun 5: 5401

Gomis-Ruth FX, Stocker W, Huber R, Zwilling R, Bode W (1993) Refined 1.8 A X-ray crystal structure of astacin, a zinc-endopeptidase from the crayfish Astacus astacus L. Structure determination, refinement, molecular structure and comparison with thermolysin. J Mol Biol 229: 945–968

Guerra D, Ballard K, Truebridge I, Vierling E (2016) S-Nitrosation of Conserved Cysteines Modulates Activity and Stability of S-Nitrosoglutathione Reductase (GSNOR). Biochemistry 55: 2452–2464

Gupta KJ, Kolbert Z, Durner J, Lindermayr C, Corpas FJ, Brouquisse R, Barroso JB, Umbreen S, Palma JM, Hancock JT, Petrivalsky M, Wendehenne D, Loake GJ (2020) Regulating the regulator: nitric oxide control of post-translational modifications. New Phytol

Hart TW (1985) Some observations concerning the S-nitroso and S-phenylsulphonyl derivatives of L-cysteine and glutathione. Tetrahedron Letters 26: 2013–2016

Hemschemeier A, Duner M, Casero D, Merchant SS, Winkler M, Happe T (2013) Hypoxic survival requires a 2-on-2 hemoglobin in a process involving nitric oxide. Proc Natl Acad Sci U S A 110: 10854–10859

Holmes MA, Matthews BW (1981) Binding of hydroxamic acid inhibitors to crystalline thermolysin suggests a pentacoordinate zinc intermediate in catalysis. Biochemistry 20: 6912–6920

Holmquist B, Vallee BL (1991) Human liver class III alcohol and glutathione dependent formaldehyde dehydrogenase are the same enzyme. Biochem Biophys Res Commun 178: 1371–1377

Huang D, Huo J, Zhang J, Wang C, Wang B, Fang H, Liao W (2019) Protein S-nitrosylation in programmed cell death in plants. Cell Mol Life Sci 76: 1877–1887

Jahnova J, Luhova L, Petrivalsky M (2019) S-Nitrosoglutathione Reductase-The Master Regulator of Protein S-Nitrosation in Plant NO Signaling. Plants (Basel) 8

Jensen DE, Belka GK, Du Bois GC (1998) S-Nitrosoglutathione is a substrate for rat alcohol dehydrogenase class III isoenzyme. Biochem J 331 (Pt 2): 659–668

Kabsch W (2010) Xds. Acta Crystallogr D Biol Crystallogr 66: 125–132

Kolbert Z, Barroso JB, Brouquisse R, Corpas FJ, Gupta KJ, Lindermayr C, Loake GJ, Palma JM, Petrivalsky M, Wendehenne D, Hancock JT (2019) A forty year journey: The generation and roles of NO in plants. Nitric Oxide 93: 53–70

Kovacs I, Holzmeister C, Wirtz M, Geerlof A, Frohlich T, Romling G, Kuruthukulangarakoola GT, Linster E, Hell R, Arnold GJ, Durner J, Lindermayr C (2016) ROS-Mediated Inhibition of S-nitrosoglutathione Reductase Contributes to the Activation of Anti-oxidative Mechanisms. Front Plant Sci 7: 1669

Kubienova L, Kopecny D, Tylichova M, Briozzo P, Skopalova J, Sebela M, Navratil M, Tache R, Luhova L, Barroso JB, Petrivalsky M (2013) Structural and functional characterization of a plant S-nitrosoglutathione reductase from Solanum lycopersicum. Biochimie 95: 889–902

Kuo EY, Chang HL, Lin ST, Lee TM (2020) High Light-Induced Nitric Oxide Production Induces Autophagy and Cell Death in Chlamydomonas reinhardtii. Front Plant Sci 11: 772

Lee U, Wie C, Fernandez BO, Feelisch M, Vierling E (2008) Modulation of nitrosative stress by S-nitrosoglutathione reductase is critical for thermotolerance and plant growth in Arabidopsis. Plant Cell 20: 786–802

Lemaire SD, Richardson JM, Goyer A, Keryer E, Lancelin JM, Makhatadze GI, Jacquot JP (2000) Primary structure determinants of the pH- and temperature-dependent aggregation of thioredoxin. Biochim Biophys Acta 1476: 311–323

Li B, Sun L, Huang J, Goschl C, Shi W, Chory J, Busch W (2019) GSNOR provides plant tolerance to iron toxicity via preventing iron-dependent nitrosative and oxidative cytotoxicity. Nat Commun 10: 3896

Lin A, Wang Y, Tang J, Xue P, Li C, Liu L, Hu B, Yang F, Loake GJ, Chu C (2012) Nitric oxide and protein S-nitrosylation are integral to hydrogen peroxide-induced leaf cell death in rice. Plant Physiol 158: 451–464

Lindermayr C (2017) Crosstalk between reactive oxygen species and nitric oxide in plants: Key role of S-nitrosoglutathione reductase. Free Radic Biol Med

Lindermayr C, Saalbach G, Durner J (2005) Proteomic identification of S-nitrosylated proteins in Arabidopsis. Plant Physiol 137: 921–930

Liu L, Hausladen A, Zeng M, Que L, Heitman J, Stamler JS (2001) A metabolic enzyme for S-nitrosothiol conserved from bacteria to humans. Nature 410: 490–494

Liu L, Yan Y, Zeng M, Zhang J, Hanes MA, Ahearn G, McMahon TJ, Dickfeld T, Marshall HE, Que LG, Stamler JS (2004) Essential roles of S-nitrosothiols in vascular homeostasis and endotoxic shock. Cell 116: 617–628

Lushchak OV, Nykorak NZ, Ohdate T, Inoue Y, Lushchak VI (2009) Inactivation of genes encoding superoxide dismutase modifies yeast response to S-nitrosoglutathione-induced stress. Biochemistry (Mosc) 74: 445–451

Marchand CH, Fermani S, Rossi J, Gurrieri L, Tedesco D, Henri J, Sparla F, Trost P, Lemaire SD, Zaffagnini M (2019) Structural and Biochemical Insights into the Reactivity of Thioredoxin h1 from Chlamydomonas reinhardtii. Antioxidants (Basel) 8

Maret W, Li Y (2009) Coordination dynamics of zinc in proteins. Chem Rev 109: 4682–4707

Matamoros MA, Cutrona MC, Wienkoop S, Begara-Morales JC, Sandal N, Orera I, Barroso JB, Stougaard J, Becana M (2020) Altered Plant and Nodule Development and Protein S-Nitrosylation in Lotus japonicus Mutants Deficient in S-Nitrosoglutathione Reductases. Plant Cell Physiol 61: 105–117

Matthews BW (1968) Solvent content of protein crystals. J Mol Biol 33: 491–497

McCall KA, Huang C, Fierke CA (2000) Function and mechanism of zinc metalloenzymes. J Nutr 130: 1437S–1446S

Merchant SS, Prochnik SE, Vallon O, Harris EH, Karpowicz SJ, Witman GB, Terry A, Salamov A, Fritz-Laylin LK, Marechal-Drouard L, Marshall WF, Qu LH, Nelson DR, Sanderfoot AA, Spalding MH, Kapitonov VV, Ren Q, Ferris P, Lindquist E, Shapiro H, Lucas SM, Grimwood J, Schmutz J, Cardol P, Cerutti H, Chanfreau G, Chen CL, Cognat V, Croft MT, Dent R, Dutcher S, Fernandez E, Fukuzawa H, Gonzalez-Ballester D, Gonzalez-Halphen D, Hallmann A, Hanikenne M, Hippler M, Inwood W, Jabbari K, Kalanon M, Kuras R, Lefebvre PA, Lemaire SD, Lobanov AV, Lohr M, Manuell A, Meier I, Mets L, Mittag M, Mittelmeier T, Moroney JV, Moseley J, Napoli C, Nedelcu AM, Niyogi K, Novoselov SV, Paulsen IT, Pazour G, Purton S, Ral JP, Riano-Pachon DM, Riekhof W, Rymarquis L, Schroda M, Stern D, Umen J, Willows R, Wilson N, Zimmer SL, Allmer J, Balk J, Bisova K, Chen CJ, Elias M, Gendler K, Hauser C, Lamb MR, Ledford H, Long JC, Minagawa J, Page MD, Pan J, Pootakham W, Roje S, Rose A, Stahlberg E, Terauchi AM, Yang P, Ball S, Bowler C, Dieckmann CL, Gladyshev VN, Green P, Jorgensen R, Mayfield S, Mueller-Roeber B, Rajamani S, Sayre RT, Brokstein P, Dubchak I, Goodstein D, Hornick L, Huang YW, Jhaveri J, Luo Y, Martinez D, Ngau WC, Otillar B, Poliakov A, Porter A, Szajkowski L, Werner G, Zhou K, Grigoriev IV, Rokhsar DS, Grossman AR (2007) The Chlamydomonas genome reveals the evolution of key animal and plant functions. Science 318: 245–250

Michelet L, Roach T, Fischer BB, Bedhomme M, Lemaire SD, Krieger-Liszkay A (2013) Down-regulation of catalase activity allows transient accumulation of a hydrogen peroxide signal in Chlamydomonas reinhardtii. Plant Cell Environ 36: 1204–1213

Morisse S, Zaffagnini M, Gao XH, Lemaire SD, Marchand CH (2014) Insight into protein S-nitrosylation in Chlamydomonas reinhardtii. Antioxid Redox Signal 21: 1271–1284

Murshudov GN, Skubak P, Lebedev AA, Pannu NS, Steiner RA, Nicholls RA, Winn MD, Long F, Vagin AA (2011) REFMAC5 for the refinement of macromolecular crystal structures. Acta Crystallogr D Biol Crystallogr 67: 355–367

Neill S, Barros R, Bright J, Desikan R, Hancock J, Harrison J, Morris P, Ribeiro D, Wilson I (2008) Nitric oxide, stomatal closure, and abiotic stress. J Exp Bot 59: 165–176

Noctor G, Mhamdi A, Chaouch S, Han Y, Neukermans J, Marquez-Garcia B, Queval G, Foyer CH (2012) Glutathione in plants: an integrated overview. Plant Cell Environ 35: 454–484

Okado-Matsumoto A, Fridovich I (2007) Putative denitrosylase activity of Cu, Zn-superoxide dismutase. Free Radic Biol Med 43: 830–836

Pace CN, Vajdos F, Fee L, Grimsley G, Gray T (1995) How to measure and predict the molar absorption coefficient of a protein. Protein Sci 4: 2411–2423

Pasquini M, Fermani S, Tedesco D, Sciabolini C, Crozet P, Naldi M, Henri J, Vothknecht U, Bertucci C, Lemaire SD, Zaffagnini M, Francia F (2017) Structural basis for the magnesium-dependent activation of transketolase from Chlamydomonas reinhardtii. Biochim Biophys Acta Gen Subj 1861: 2132–2145

Pérez-Pérez ME, Mauriès A, Maes A, Tourasse NJ, Hamon M, Lemaire SD, Marchand CH (2017) The Deep Thioredoxome in Chlamydomonas reinhardtii: New Insights into Redox Regulation. Mol Plant 10: 1107–1125

Pokora W, Aksmann A, Bascik-Remisiewicz A, Dettlaff-Pokora A, Rykaczewski M, Gappa M, Tukaj Z (2017) Changes in nitric oxide/hydrogen peroxide content and cell cycle progression: Study with synchronized cultures of green alga Chlamydomonas reinhardtii. J Plant Physiol 208: 84–93

Ren X, Sengupta R, Lu J, Lundberg JO, Holmgren A (2019) Characterization of mammalian glutaredoxin isoforms as S-denitrosylases. FEBS Lett 593: 1799–1806

Robert X, Gouet P (2014) Deciphering key features in protein structures with the new ENDscript server. Nucleic Acids Res 42: W320–324

Rouhier N, Lemaire SD, Jacquot JP (2008) The role of glutathione in photosynthetic organisms: emerging functions for glutaredoxins and glutathionylation. Annu Rev Plant Biol 59: 143–166

Sakamoto A, Ueda M, Morikawa H (2002) Arabidopsis glutathione-dependent formaldehyde dehydrogenase is an S-nitrosoglutathione reductase. FEBS Lett 515: 20–24

Sanghani PC, Bosron WF, Hurley TD (2002) Human glutathione-dependent formaldehyde dehydrogenase. Structural changes associated with ternary complex formation. Biochemistry 41: 15189–15194

Sanghani PC, Davis WI, Zhai L, Robinson H (2006) Structure-function relationships in human glutathione-dependent formaldehyde dehydrogenase. Role of Glu-67 and Arg-368 in the catalytic mechanism. Biochemistry 45: 4819–4830

Sanz-Luque E, Ocana-Calahorro F, Llamas A, Galvan A, Fernandez E (2013) Nitric oxide controls nitrate and ammonium assimilation in Chlamydomonas reinhardtii. J Exp Bot 64: 3373–3383

Scaife MA, Nguyen G, Rico J, Lambert D, Helliwell KE, Smith AG (2015) Establishing Chlamydomonas reinhardtii as an industrial biotechnology host. Plant J 82: 532–546

Scaife MA, Smith AG (2016) Towards developing algal synthetic biology. Biochem Soc Trans 44: 716–722

Sengupta R, Holmgren A (2012) The role of thioredoxin in the regulation of cellular processes by S-nitrosylation. Biochim Biophys Acta 1820: 689–700

Shao N, Beck CF, Lemaire SD, Krieger-Liszkay A (2008) Photosynthetic electron flow affects H(2)O (2) signaling by inactivation of catalase in Chlamydomonas reinhardtii. Planta 228: 1055–1066

Shao Z, Borde C, Marchand CH, Lemaire SD, Busson P, Gozlan JM, Escargueil A, Marechal V (2019) Detection of IgG directed against a recombinant form of Epstein-Barr virus BALF0/1 protein in patients with nasopharyngeal carcinoma. Protein Expr Purif 162: 44–50

Sievers F, Wilm A, Dineen D, Gibson TJ, Karplus K, Li W, Lopez R, McWilliam H, Remmert M, Soding J, Thompson JD, Higgins DG (2011) Fast, scalable generation of high-quality protein multiple sequence alignments using Clustal Omega. Mol Syst Biol 7: 539

Skelly MJ, Malik SI, Le Bihan T, Bo Y, Jiang J, Spoel SH, Loake GJ (2019) A role for S-nitrosylation of the SUMO-conjugating enzyme SCE1 in plant immunity. Proc Natl Acad Sci U S A 116: 17090–17095

Sreerama N, Woody RW (2000) Estimation of protein secondary structure from circular dichroism spectra: comparison of CONTIN, SELCON, and CDSSTR methods with an expanded reference set. Anal Biochem 287: 252–260

Stomberski CT, Anand P, Venetos NM, Hausladen A, Zhou HL, Premont RT, Stamler JS (2019) AKR1A1 is a novel mammalian S-nitroso-glutathione reductase. J Biol Chem 294: 18285–18293

Stomberski CT, Hess DT, Stamler JS (2019) Protein S-Nitrosylation: Determinants of Specificity and Enzymatic Regulation of S-Nitrosothiol-Based Signaling. Antioxid Redox Signal 30: 1331–1351

Ticha T, Cincalova L, Kopecny D, Sedlarova M, Kopecna M, Luhova L, Petrivalsky M (2017) Characterization of S-nitrosoglutathione reductase from Brassica and Lactuca spp. and its modulation during plant development. Nitric Oxide 68: 68–76

Ticha T, Lochman J, Cincalova L, Luhova L, Petrivalsky M (2017) Redox regulation of plant S-nitrosoglutathione reductase activity through post-translational modifications of cysteine residues. Biochem Biophys Res Commun 494: 27–33

Umbreen S, Lubega J, Cui B, Pan Q, Jiang J, Loake GJ (2018) Specificity in nitric oxide signalling. J Exp Bot 69: 3439–3448

Vagin A, Teplyakov A (2010) Molecular replacement with MOLREP. Acta Crystallogr D Biol Crystallogr 66: 22–25

van Stokkum IH, Spoelder HJ, Bloemendal M, van Grondelle R, Groen FC (1990) Estimation of protein secondary structure and error analysis from circular dichroism spectra. Anal Biochem 191: 110–118

Vavitsas K, Crozet P, Vinde MH, Davies F, Lemaire SD, Vickers CE (2019) The Synthetic Biology Toolkit for Photosynthetic Microorganisms. Plant Physiol 181: 14–27

Wallace AC, Laskowski RA, Thornton JM (1995) LIGPLOT: a program to generate schematic diagrams of protein-ligand interactions. Protein Eng 8: 127–134

Wei L, Derrien B, Gautier A, Houille-Vernes L, Boulouis A, Saint-Marcoux D, Malnoe A, Rappaport F, de Vitry C, Vallon O, Choquet Y, Wollman FA (2014) Nitric oxide-triggered remodeling of chloroplast bioenergetics and thylakoid proteins upon nitrogen starvation in Chlamydomonas reinhardtii. Plant Cell 26: 353–372

Whitmore L, Wallace BA (2004) DICHROWEB, an online server for protein secondary structure analyses from circular dichroism spectroscopic data. Nucleic Acids Res 32: W668–673

Wijffels RH, Kruse O, Hellingwerf KJ (2013) Potential of industrial biotechnology with cyanobacteria and eukaryotic microalgae. Curr Opin Biotechnol 24: 405–413

Wilson DK, Quiocho FA (1993) A pre-transition-state mimic of an enzyme: X-ray structure of adenosine deaminase with bound 1-deazaadenosine and zinc-activated water. Biochemistry 32: 1689–1694

Xu S, Guerra D, Lee U, Vierling E (2013) S-nitrosoglutathione reductases are low-copy number, cysteine-rich proteins in plants that control multiple developmental and defense responses in Arabidopsis. Front Plant Sci 4: 430

Yu M, Lamattina L, Spoel SH, Loake GJ (2014) Nitric oxide function in plant biology: a redox cue in deconvolution. New Phytol 202: 1142–1156

Yu Z, Cao J, Zhu S, Zhang L, Peng Y, Shi J (2020) Exogenous Nitric Oxide Enhances Disease Resistance by Nitrosylation and Inhibition of S-Nitrosoglutathione Reductase in Peach Fruit. Front Plant Sci 11: 543

Zaffagnini M, Bedhomme M, Groni H, Marchand CH, Puppo C, Gontero B, Cassier-Chauvat C, Decottignies P, Lemaire SD (2012) Glutathionylation in the photosynthetic model organism Chlamydomonas reinhardtii: a proteomic survey. Mol Cell Proteomics 11: M111014142

Zaffagnini M, De Mia M, Morisse S, Di Giacinto N, Marchand CH, Maes A, Lemaire SD, Trost P (2016) Protein S-nitrosylation in photosynthetic organisms: A comprehensive overview with future perspectives. Biochim Biophys Acta 1864: 952–966

Zaffagnini M, Fermani S, Marchand CH, Costa A, Sparla F, Rouhier N, Geigenberger P, Lemaire SD, Trost P (2019) Redox Homeostasis in Photosynthetic Organisms: Novel and Established Thiol-Based Molecular Mechanisms. Antioxid Redox Signal 31: 155–210

Zaffagnini M, Marchand CH, Malferrari M, Murail S, Bonacchi S, Genovese D, Montalti M, Venturoli G, Falini G, Baaden M, Lemaire SD, Fermani S, Trost P (2019) Glutathionylation primes soluble glyceraldehyde-3-phosphate dehydrogenase for late collapse into insoluble aggregates. Proc Natl Acad Sci U S A 116: 26057–26065

Zaffagnini M, Michelet L, Sciabolini C, Di Giacinto N, Morisse S, Marchand CH, Trost P, Fermani S, Lemaire SD (2014) High-resolution crystal structure and redox properties of chloroplastic triosephosphate isomerase from Chlamydomonas reinhardtii. Mol Plant 7: 101–120

Zaffagnini M, Morisse S, Bedhomme M, Marchand CH, Festa M, Rouhier N, Lemaire SD, Trost P (2013) Mechanisms of nitrosylation and denitrosylation of cytoplasmic glyceraldehyde-3-phosphate dehydrogenase from Arabidopsis thaliana. J Biol Chem 288: 22777–22789

Zalutskaya Z, Kochemasova L, Ermilova E (2018) Dual positive and negative control of Chlamydomonas PII signal transduction protein expression by nitrate/nitrite and NO via the components of nitric oxide cycle. BMC Plant Biol 18: 305

Zhan N, Wang C, Chen L, Yang H, Feng J, Gong X, Ren B, Wu R, Mu J, Li Y, Liu Z, Zhou Y, Peng J, Wang K, Huang X, Xiao S, Zuo J (2018) S-Nitrosylation Targets GSNO Reductase for Selective Autophagy during Hypoxia Responses in Plants. Mol Cell 71: 142–154 e146

Zhang T, Ma M, Chen T, Zhang L, Fan L, Zhang W, Wei B, Li S, Xuan W, Noctor G, Han Y (2020) Glutathione-dependent denitrosation of GSNOR1 promotes oxidative signalling downstream of H2 O2. Plant Cell Environ

